# BuzzWatch: Uncovering Multi-scale Temporal Patterns in Mosquito Behavior Through Continuous Long-term Monitoring

**DOI:** 10.1101/2025.01.24.634688

**Authors:** Théo Maire, Zhong Wan, Louis Lambrechts, Felix J.H. Hol

**Author notes:** **For correspondence:** (FJHH).

## Abstract

Understanding the temporal dynamics of mosquito behavior is essential for developing effective interventions against pathogen transmission. However, limited knowledge exists about the environmental, physiological, and genetic factors influencing mosquito activity patterns. This knowledge gap is partly due to a lack of tools to accurately quantify the behavior of free-flying mosquitoes over extended periods. Here, we introduce BuzzWatch, an open-source, low-cost platform designed to continuously monitor mosquito flight behavior over several weeks with high temporal resolution. BuzzWatch records videos of mosquitoes freely flying in a transparent cage and automates the extraction, analysis, and visualization of behavioral data, including flight trajectories and population-level flight and sugar-feeding statistics. Using BuzzWatch, we quantified the daily rhythms of 10 *Aedes aegypti* populations from various geographic origins. Globally invasive Ae. aegypti showed increased sugar feeding and flight activity during midday compared to native African populations. Our platform further revealed subtle, long-lasting effects of blood feeding on activity patterns and a complex response to extended daylight periods. By integrating a host-seeking module in BuzzWatch to deliver CO_2_ and heat pulses, we observed a twofold increase in *Ae. aegypti*’s response to host-associated cues during the daytime compared to nighttime. Combined, these results demonstrate BuzzWatch’s potential to investigate responses to host cues over seconds, natural variability in daily rhythms over hours, and phenotypic plasticity over days. BuzzWatch offers a novel perspective on mosquito behavior over multiple timescales, paving the way for advanced ecological and epidemiological studies that can inform targeted and effective vector control strategies.

**Short Summary:** This study introduces a new experimental platform to monitor multiple features of mosquito behavior (flight activity, sugar feeding, short-range host-seeking) with high temporal resolution over several weeks.

**Highlights:**

- Continuous month-long tracking of mosquito flight activity in a laboratory setting.
- Multiscale analysis of flight behavior, from seconds to weeks.
- Mapping of natural variation in daily rhythm of *Aedes aegypti* populations.
- Quantifying the long-lasting effect of physiological (blood-meal digestion) and environmental (photoperiod increase) perturbations.
- Automatically monitoring response to short-range host-seeking cues at specific times of the day.

## INTRODUCTION

In the fleeting weeks of their lifespan, female mosquitoes engage in a sequence of crucial behaviors—mating, blood-feeding, and egg-laying—to ensure successful reproduction. Marked progress has been made regarding the specific cues and associated neurosensory systems that allow mosquitoes to precisely locate a host (reviewed in Coutinho-Abreu, Riffell, and Akbari 2022) or an oviposition site (Afify and Galizia 2015), yet we have a limited understanding of the factors that dictate the timing of these behaviors. Mosquito do not engage in specific behaviors uniformly over time, yet behaviors rather concentrate at specific periods of the day—usually before sunset for *Aedes* species (Zahid et al. 2023) and after sunset for *Anopheles* (Kenea et al. 2016) and *Culex* species (Duffield 2024). The timing of blood feeding, for instance, is typically estimated using the “Human Landing Catch” assay, in which an individual records mosquito landings on their own body over time (for example in (Captain-Esoah et al. 2020). Although this field method accounts for relevant environmental variations and human cues, it is very labor intensive and complicates the isolation of specific factors that determine timing of host-seeking behavior. Laboratory experiments conducted in the absence of human cues but with periodic light cycles have shown that mosquitoes display rhythmic activity patterns (M. D. R. Jones, Hill, and Hope 1967; Taylor and M. D. R. Jones 1969), mirroring observations from field studies. Like in other insects, these behaviors are governed by an internal circadian rhythm, modulated by environmental cues like sunlight, but can also persist without them. At the molecular level, circadian rhythms in mosquitoes are regulated by clock genes, such as *period* and *timeless*, which control proteins driving daily activity cycles (Duffield 2024).

Historically, the characterization of circadian rhythms in mosquitoes was made possible by remarkably simple behavioral assays. These methods included using microphones to record the flight activity of individual mosquitoes (M. Jones 1964) and infrared beams to measure walking activity in narrow tubes (e.g. Newman, Anderson, and Goldberg 2016; Ajayi et al. 2024). While these assays have provided valuable insights, they are inadequate for comprehensive flight activity monitoring. Typically, measurements span only a few hours to days and focus on basic indicators like “active” or “not active,” and, importantly, severely restrict mosquito movement. Consequently, these methods do not allow quantification of rhythmic behaviors involving flight, such as sugar feeding, host-seeking, or egg-laying.

While assays to longitudinally monitor the activity of mosquitoes are limited, various recent studies have used high-resolution video recording with multiple cameras to track 3D mosquito flight. In conjunction with machine vision techniques, these approaches typically allow for detailed analysis of flight-based behaviors such as swarming (Cavagna et al. 2023), escape from predators (Cribellier et al. 2024), or navigating toward human cues (van Breugel et al. 2015). However, these setups are costly and generate vast amounts of data (Spitzen and Takken 2018), making them suitable for short-term (typically several minutes), high-resolution monitoring but inappropriate for long-term tracking over weeks. To date, no setup has been optimized to monitor the activity of freely flying of mosquitos over long periods of time.

Nevertheless, robustly recording and analyzing mosquito flight activity over several weeks is meaningful for several reasons: it allows measuring the effects of physiological processes like pathogen infection or blood ingestion that have long-lasting impacts, and it establishes a foundation to examine the complex interplay between flight activity and specific behaviors such as host-seeking or oviposition. Furthermore, long-term monitoring could enable comparative studies to better understand genetic factors driving variability between closely related species or strains. Additionally, this approach could be valuable for studying and quantifying behavioral adaptations in response to public health interventions such as insecticide treatments. With these goals and applications in mind, we sought to construct a low-cost open-source setup to monitor the flight activity of mosquitoes over weeks and enable quantification of specific behaviors like sugar-feeding and host-seeking.

Here we report “BuzzWatch”, an open-source hardware platform and computational pipeline to measure and analyze the rhythmic behavior of mosquitoes over weeks. Alongside this manuscript, we offer tutorials on constructing the setup and analyzing the data on a dedicated wiki page, and a graphical user interface to facilitate mosquito tracking from video footage and to perform multi-scale analysis from mosquito tracks. These resources will ensure reproducibility and promote consistent data exploration across different experiments and laboratories. In this work, we focus on *Aedes aegypti*, the primary vector of many arboviruses, and present four independent case studies, that collectively demonstrate the extensive capabilities of BuzzWatch:

1. Mapping natural variation in the daily rhythm of *Aedes aegypti* colonies, revealing a distinct signature distinguishing native and invasive populations.
2. Quantifying the long-term impact of physiological (ingestion of a blood-meal) or environmental changes (modification of the light cycle) on mosquito activity patterns, highlighting aspects of their phenotypic plasticity.
3. Measuring responses to repeated pulses of host cues, uncovering consistent differences between responses during night and day times.

In this work, we use the term ‘daily rhythm’ as an umbrella term to describe the observable patterns of mosquito behavior over a 24-hour period, acknowledging that these patterns may include both internal circadian influences and external environmental factors.

## RESULTS

### An open platform to continuously monitor mosquito flight activity over weeks

To monitor mosquito behavior over weeks, we designed a dedicated hardware and software platform, constructed from readily available and low-cost materials and open-source methods to monitor the activity of populations of 40-60 adult mosquitoes (Fig.1A).

**Figure 1.**
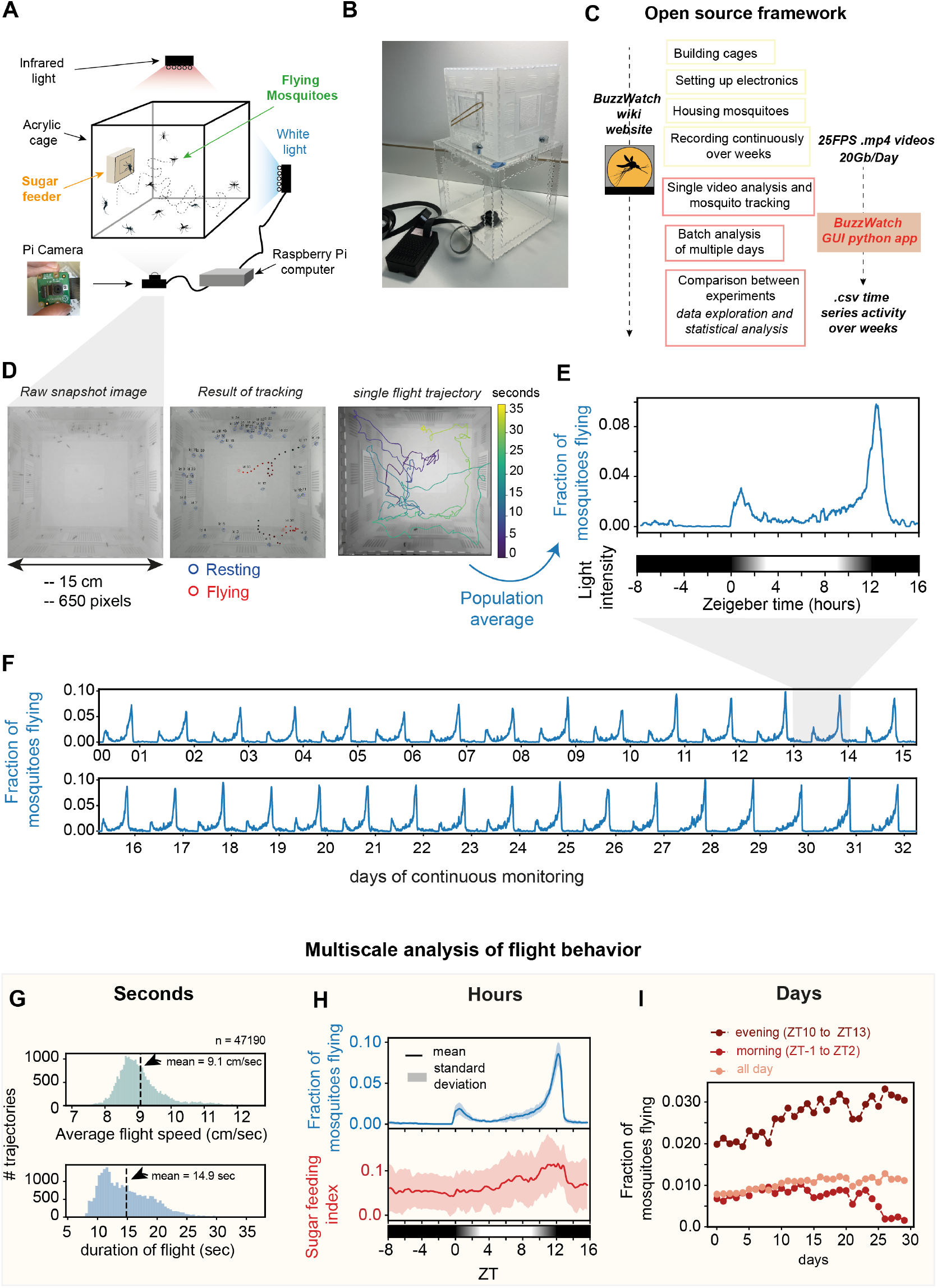
BuzzWatch: an open platform for long-term mosquito flight activity monitoring. **A**. Cartoon of the set-up with the minimal set of elements required to monitor mosquito flight activity over weeks. **B**. Pictures of transparent acrylic cage with Raspberry Pi and Camera NoIR V2 on the bottom. See Supp. Fig1 for details and assembly instructions. **C**. Flow chart of BuzzWatch pipeline for constructing the complete set-up and analyzing video data. Step-by-step guides are available in a dedicated website (https://theomaire.github.io/buzzwatch/). A GUI python app (https://github.com/theomaire/BuzzWatch_app_v2) facilitates analysis from raw .mp4 video files to comparisons between experiments, and statistical analysis (see Fig 2-4.) **D**. Left: Raw snapshot image from a typical .mp4 movie as registered by the Pi Camera. Center: Overlay of the image with identity tracker, blue circles correspond to resting mosquitos, red circles indicate flying mosquitoes. Filled black to red circles indicate position of flying mosquito in the previous 20 frames. Right: Overlay of camera image showing a 37 second-long flight trajectory of a single mosquito. Line color indicates time. **E**. Top: Graph showing the instantaneous fraction of flying mosquitoes of a cohort of 40 *Ae. aegypti* females for 24 hours (moving average over 20 minutes). Bottom: Horizontal bar showing light intensity profile inside the set-up over a 24 hour cycle. We use the Zeitgeber time notation and the light cycle is programmed to have 12 hours of white light gradually changing (from ZT0 to ZT12), and12 hours of complete darkness (ZT12 to ZT24) and mimicking real sunlight variation. By convention we use a periodic boundary notation for ZT times before ZT0, such that e.g. ZT-8 is the same as ZT16. **F**. Graph showing the instantaneous fraction of flying mosquitoes of a cohort of 40 *Ae. aegypti* females for 31 days (moving average over 60 minutes). **G** Statistics of flight trajectories from the same experiment as 1E. Histograms showing the average flight speed (green bars, top panel) and the duration of flight (blue bars, bottom panel) of *n* = 47, 190 flight trajectories. For flight speed (top graph), the average speed is computed from first averaging instantaneous speed over single trajectories and then averaging over all trajectories. As we use 2D trajectories, the average value of 9.1cm/sec does not correspond to the “real” flight speed, but is rather a 2D approximation. Assuming a uniform distribution over the height of the cage and isotropic movement, the 3D flight speed can be approximated by applying a factor 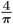 to the speed of the 2D projection (see Supplementary text 1), yielding 11.5cm/sec. **H**. Statistics over a day. Top, blue line indicates the instantaneous fraction of flying mosquitoes averaged over 32 days. Bottom, red line indicates sugar feeding index (as defined in Supp. Fig.S3 A-D) dashed lines are +/− std of days. **I**. Statistics over a month. Dotted lines indicate average fraction flying averaged over a given day interval, for each day of the experiment.

#### Hardware: continuous monitoring of flight activity

We constructed a 15 × 15 × 15 cm transparent cage to house mosquitoes, allowing them to rest, fly, and sugar feed at any time (Fig.1B). The cage includes a sugar feeder supporting a population of 40 to 60 mosquitoes to thrive without human intervention for weeks, provided that the cage is placed in an environment with appropriate temperature and humidity conditions. The cage, sugar feeder, and supporting structural elements are constructed out of acrylic using a laser cutter, facilitating easy reproduction and modification. We designed and tested several versions of the setup, which can be placed in either a climatic room or a climatic controlled environment where temperature, humidity, and lighting are externally controlled (Fig.S1E,F), or in a custom-built environment where artificial daylight intensity is directly controlled by an RGB LED array connected to a Raspberry Pi (Fig.S1C,D).

We use the Raspberry Pi single board computer and associated Pi Camera mounted beneath the cage as a cost-effective means to record flight activity for weeks (Fig.1B, Fig.S1). Infrared illumination and a long-pass filter facilitate video acquisition independent of environmental lighting conditions without impacting mosquito behavior (infrared wavelengths do provide limited cues to mosquitoes (Zermoglio et al. 2017; Muir, Thorne, and Kay 1992). In balancing the quality of video acquisition required for the detection and tracking of flying mosquitoes with the reliability and storage capacity over weeks, we chose to record at a “low” resolution of 650×650 pixels at a relatively high frame rate of 25 frames per second. This allows all mosquitoes within the cage to be visible and enables reliable tracking of flying mosquitoes (Fig.1D). Recordings result in week- or month-long movies (segmented into 20-minute videos) of 40-60 freely behaving mosquitoes in a cage.

The list of parts and technical information necessary to construct the set-up are briefly described in the Methods section (Table 2) and extensively documented in a dedicated “wiki” website with step-by-step illustrated guides (Fig.1C, https://theomaire.github.io/buzzwatch/construct.html). The total cost of materials for one complete set-up is around 150-300euros per cage (see Table 2)), making it affordable for laboratories to build several replicas.

**Table 1.**
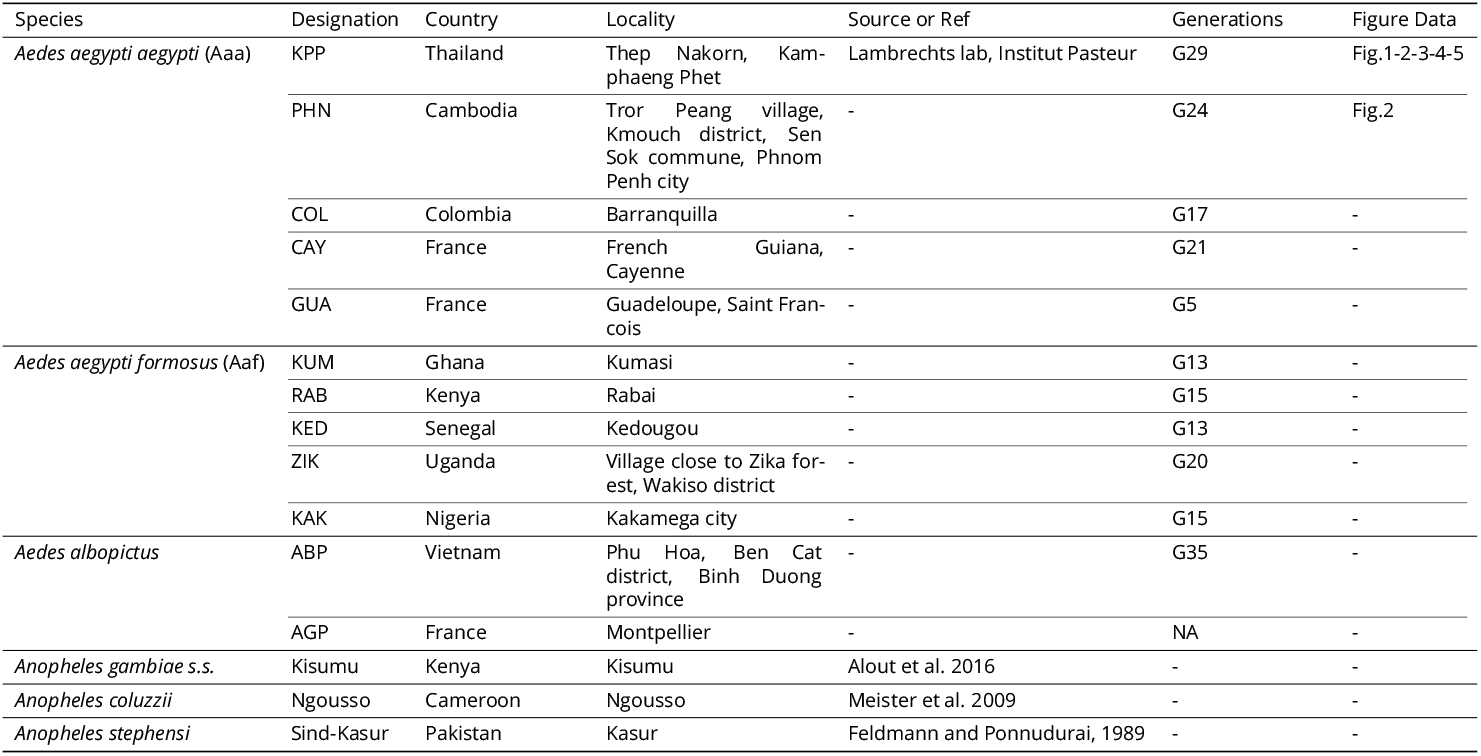
Mosquito strains used in this study.

**Table 2.** BuzzWatch parts list.

| Part Name | Specific Reference | Indicative Price |
| --- | --- | --- |
| **BuzzWatch Cage** |  |  |
| Acrylic sheets 3mm for cage | Cast PPMA transparent and diffusive from "Richardson" | 10 / cage (700mm x 400mm sheet) |
| Acrylic sheets 1.5mm for windows | Cast PPMA transparent from Richardson |  |
| Micro computer | Raspberry Pi 4B+ 4G RAM | 65 |
| Micro-SD card | 64GB microSD card. SanDisk Ultra A1 MicroSDXC | 15 |
| Camera | Raspberry Pi NoIR Camera Board v2 - 8 Megapixels | 25-30 |
| SSD | 512GB SSD external hard drive - (Samsung) Portable SSD T7 | 100 |
| Infrared filter | Long Pass Filter 850nm, 25mm diameter FGL850M (Thorlabs) | 60 |
| Case camera | Adafruit Raspberry Pi P3253 (gotronic) | 3 |
| Camera connector | RB-CAM-1000 35861 Gotronic (1m version) | 5 |
| Case | Rpi 4 case Joy-it. CASEP4+3B 36489 | 15 |
| Power supply | USB-C Power supply 5.1V 3A (official Raspberry Pi 4 power supply) | 8-10 |
| Infrared illumination | Andoer IR49S INFRARED | 25 |
| **Environmental Container** |  |  |
| White light illumination | Neopixels 60 LEDs RGB 2 meters (Adafruit) | 25-30 |
| Temperature humidity sensor | DHT11 ST052 | 3 |
| Light sensor | ADA4698 | 18 |
| Jump cables wires | Any |  |
| Acrylic sheets | White opaque acrylic 3mm PPMA from Richardson |  |
| Aluminum rail for structure | "Profilé Aluminium RS PRO 30 x 30 mm x 2000mm" | 20-30/meter |
| Screws<br>12V DC fans | Any<br>be quiet! Pure Wings 2 80 mm PWM | 15 |
| **Host-seeking Module** |  |  |
| CO<sub>2</sub> With pressure regulator | TROPICA Plant Growth System Nano - with 95g CO<sub>2</sub> canister | 90 |
| Air pump | Air pump 5V, ZR370-02PM | 8 |
| Silicon air tubes | Any | 5-10 |
| Module Peltier | Module Peltier 5 Vcc TEC1-04905 | 8 |
| Solenoid valve | U.S. Solid 12V DC 3/4" G Electric Solenoid Valve Brass Normally Closed "NC" | 45 |
| 5V relay module | Yizhet 5V Relay Module DC 5V 230V 8 Channel | 10-15 |
| Power supply 5V and 12V for Peltier, solenoid, pump | Any | 5-10 per power supply |
| CO<sub>2</sub> sensor | CO<sub>2</sub> SEN0159 (DF robot) or ENS160 ADA5606 | 65 or 30 |
| Connector Jack DC female 12V | Any | 1-5 |

#### Software: Tracking and measuring mosquito flight activity

We developed a dedicated Python pipeline to extract and analyze the flight activity of mosquitoes captured during month-long recordings, which can be used as a command line tool or through a dedicated graphical user interface facilitating detailed analysis (Fig.1C). With its straightforward design and virtually no user input required, our pipeline using parallel computing typically processes 20-minute videos in 1-5 minutes, enabling the analysis of one week’s data in 10 hours on a standard laptop computer.

Leveraging the observation that every alive mosquito moves at least once per day, we create background images devoid of alive mosquitoes by averaging 100 frames over a span of 33 hours (see Fig.S2). Averaged images facilitate daily counting of the number of dead mosquitos to estimate the number of alive mosquitos at any time over a 30-day experiment. We employed computer vision methods to detect centroids of mosquitoes (Fig.1D middle, Fig.S2A).

We refined the greedy centroid tracking algorithm to efficiently classify individual mosquitoes as resting or flying. This classification allows robust quantification of population level activity patterns and metrics such as the fraction of flying mosquitoes at any given time (Fig.1E) and observe the longitudinal dynamics of rhythmic patterns over long timescales (Fig.1F). We furthermore extract the 2D projection of the 3D flight trajectories of mosquitoes (Fig.1D right, Fig.S2B,C,D,E). While 2D projection underestimates flight speed by a factor of 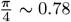 (assuming isotropic movement in a 3 dimensions, see Supplementary text1) this approximation does not affect the quantification of population level activity patterns and metrics such as the fraction of flying mosquitoes or the duration of individual flight bouts. Describing mosquito flight behavior in detail is complex, yet simply knowing when a mosquito starts or stops flying can provide significant biological insights as initiating flight is typically the first step towards behaviors such as sugar feeding or host seeking.

In Fig.1D-I, we present an example dataset for the continuous 32-day monitoring of 40 non-blood-fed Ae. aegypti females. For this experiment (and all other if not specified otherwise), we used a 24-hour cycle consisting of 12 hours of light and 12 hours of darkness, with a smooth dimming transition between the light and dark. According to the Zeitgeber convention, ZT0 marks the beginning of the increase in light intensity (artificial ‘sunrise’), and ZT12 marks the end of the decrease in light intensity before complete darkness (artificial ‘sunset’).

Consistent with previous studies on Ae. aegypti locomotor and flight activity (Taylor and M. D. R. Jones 1969), we observed two peaks of flight activity: a smaller peak following “sunrise” (ZT0) and a more pronounced peak before “sunset” (ZT12) (Fig.1E). During the night and midday flight activity is low and Ae. aegypti females are generally resting. At the peak of activity, only about 10% of mosquitoes are flying at any moment, while nearly all fly at least once during the evening peak (Fig.S4). We focus on the instantaneous fraction flying, as it offers a robust measure indicative of the transition between resting and flying. In addition to the fraction of flying mosquitoes, a population measure, we also quantified several features of individual flight trajectories, including the duration of flight, the duration of resting, and the 2D projection of instantaneous/average flight speed (Fig.S2; Fig.S3). By monitoring a group of 40 mosquitoes for several weeks, an experiment typically yields around 10,000 flight trajectories, which form a solid basis to compute robust distributions of the different flight statistics (Fig.1G).

#### Multiscale analysis from seconds to weeks

With datasets capturing flight activity at sub-second resolution over weeks, we were able to perform nuanced statistical analyses across multiple time scales—from mere seconds to entire days (Fig.1G-I). These rich data facilitate the examination of three key forms of variability: between individual flight trajectories (Fig.1G); variation during the day (24 hours), quantified by averaging the data over multiple days to discern daily rhythmic patterns and day-to-day variations in mosquito activity (Fig.1H); and across consecutive days (or parts of the day) to understand longitudinal trends over weeks (Fig.1I). For the fraction of flying *Ae. aegypti*, we observed that the daily activity pattern remained remarkably similar over 32 days (Fig.1H top). However, we observed an increase in peak activity in the evening over the course of the experiment, while the morning peak started to decrease after 20 days. This dual pattern of consistent daily rhythms coupled with gradual changes over weeks portrays the flight activity rhythm of mosquito as a complex behavioral trait, which emerges from the interplay between robust internal regulation and nuanced physiological changes.

Next, we repeated the multi-scale analysis on different behaviors or flight features such as sugar-feeding activity, flight duration, and flight speed. To estimate sugar-feeding behavior, we compared mosquito presence on the sugar-feeder to that on a control area, establishing a “sugar feeding index” (Fig.S3A-D). This method revealed increased presence on the sugar-feeder around the evening peak (ZT10-ZT12), suggesting increased sugar feeding (Fig.S3D). Compared to the instantaneous fraction flying, we observed a less pronounced pattern for sugar feeding index throughout the day, significant variation between days, and no discernable trend over long time-scales (Fig.S3E). Similarly, for flight duration and speed, no distinct daily or long-term pattern emerged (Fig.S3F-G), indicating high stability of these behavioral features in the absence of external perturbation.

The data presented above demonstrate that BuzzWatch provides a robust and efficient way to quantify the multiscale temporal dynamics of mosquito flight activity patterns over weeks. Using the dedicated python GUI app and guidelines from the wiki, all analysis and plots can be reproduced with a few clicks. Having established the method using *Ae. aegypti* maintained in a stable environment and isolated from host cues, we next sought to uncover the relative contribution of genetic, physiological, and environmental factors to the daily flight activity rhythm of mosquitoes.

### BuzzWatch uncovers intra-species variation in the daily rhythm of flight activity

Having demonstrated our method using a single *Ae. aegypti* mosquito cohort, we sought to probe the range of natural variability mosquito may display, from species to sub-species, colonies, and individual replicate populations.

#### Monitoring nocturnal and diurnal species

We first compared the activity patterns of *Ae. aegypti* to a closely related mosquito species, *Aedes albopictus* and a more distant species *Anopheles stephensi* (Fig.2A). As expected, we observed major differences in both the timing and amplitude of activity throughout the day, *An. stephensi* was mostly active during the night, whereas *Ae. aegypti* and *Ae. albopictus* were completely inactive at night. On the other hand, closely related species like *Ae. aegypti* and *Ae. albopictus* have a globally similar timing of flight activity, yet *Ae. albopictus* showed much higher total activity. These results demonstrate that our platform is compatible with a large variety of different mosquito species and recovers the known difference between key species.

**Figure 2.**
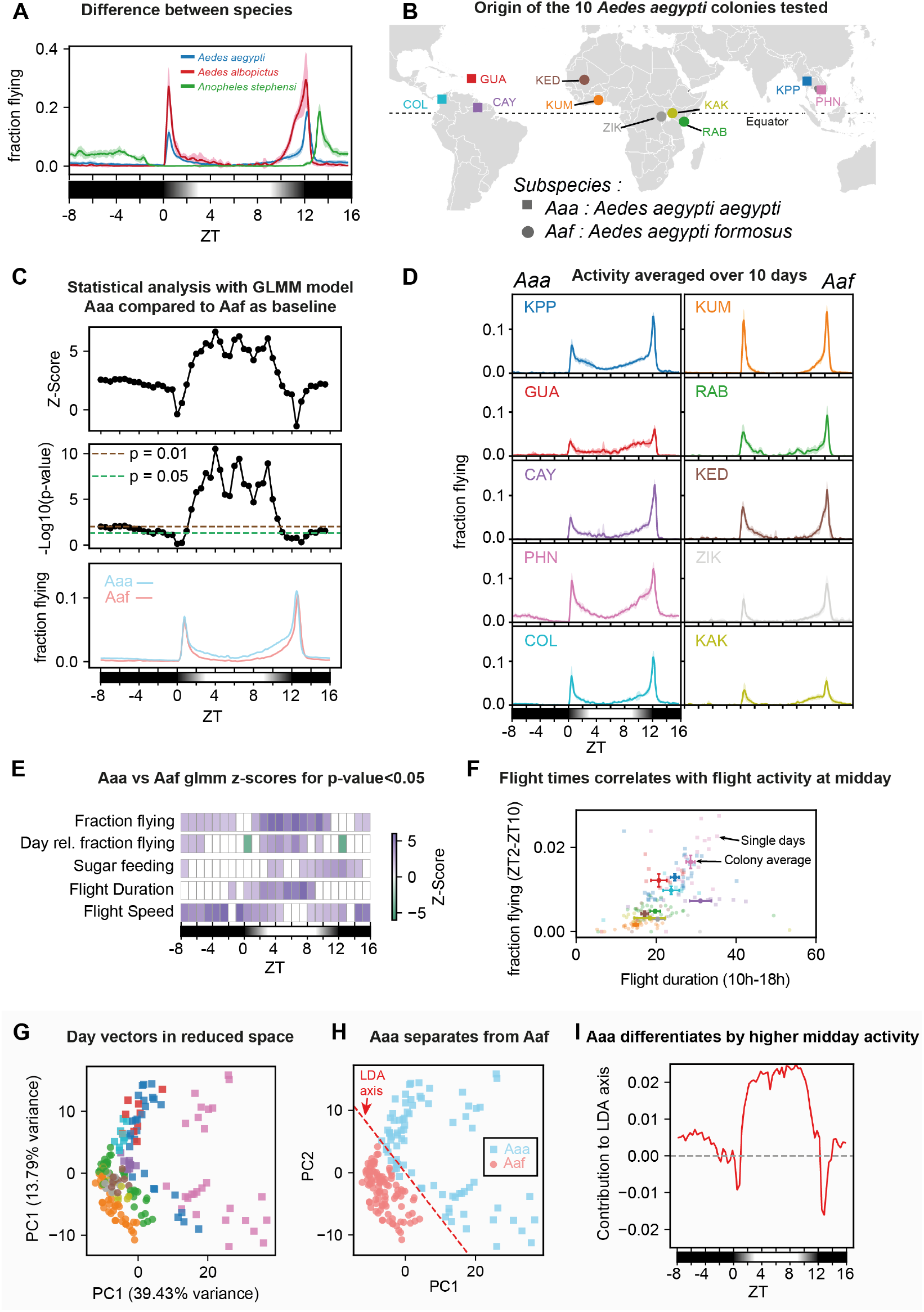
BuzzWatch uncovers distinct flight behavior patterns of *Aedes aegypti* subspecies. **A**. Fraction of mosquitoes flying, 20min moving average, dashed lines are std over 10 consecutive days. Aedes albopictus ABP, Aedes aegypti KPP, Anopheles stephensi (Sind-Kasur Nijmegen strain (Feldmann and Ponnudurai 1989)). **B**. Map of the initial sampling location of *Aedes aegypti* colonies, circles (for *Aaa*) and squares (for *Aaf*) represent the original sampling location.**C**. Generalized linear mixed model (GLMM) statistical test on fraction flying at different time of the day (1 hour interval) between *Aaa* and *Aaf* groups (each composed of 5 colonies monitored for 10 days with 1-3 replicates). Different days are interpreted as independent variables. Independent tests were performed for each hour of the day. **(Top)** z-score for each test as function of time of the day. **(Middle)** p-value associated with the z-score. **(Bottom)** 20 min moving average, averaged over 10 days for each colony. Blue line shows the average for *Aaa* colonies (KPP-PHN-GUA-COL-CAY), and red line the average of *Aaf* colonies (RAB-KUM-KED-KAK-ZIK). **D**. Activity profiles averaged over 10 days for each colony, dashed lines are std over 10 consecutive days. **E**. Summary heatmap of GLMM test similar to **C** (average over 1-hour bin, *Aaa* vs *Aaf*) for 6 different variables: “fraction flying” is the instantaneous fraction flying as in 1; “Day Rel. fraction flying” is the instantaneous fraction flying normalized to the cumulative sum of fraction flying during a day (day-relative); “Sugar feeding” is the difference between the fraction of mosquito resting on the sugar feeder and the fraction of mosquito on a surface of similar dimension without sugar feeder; “Flight duration” is the average duration of flight tracks in seconds; “Flight speed” is the average speed of flight tracks in cm/second. Color indicates the value of z-scores (purple for positive, green for negative), only z-scores corresponding to a p-value < 0.05 are displayed. **F**. Scatter plot of flight duration vs. fraction flying for the period ZT2-ZT10. Each dot corresponds to the value of a single day of a single cage of a single colony. Color code for the colonies as in B-D. Pearson correlation coefficient *ρ* = 0.51, p-value <0.001. Dots with error bar correspond to average over all the data points from a given colony, error bar is standard deviation. **G**. PCA dimension reduction, 1 point represents 1 day of 1 replicate of 1 colony. *Aaf* are circles and *Aaf* are squares. Data used for dimensionality reduction are (mean; std; min; max) of the fraction flying computed in 20 minute bins. **H**. Linear dispersion analysis over the PCA-reduced coordinates (PC1 and PC2 only) to find the direction (dashed red line) separating *Aaa* (blue) from *Aaf* (red) points. **I**. Relative contribution of features from specific time intervals to the direction separating *Aaa* from *Aaf* best in PCA-reduced space (H).

#### Mapping the behavior of 10 geographically diverse *Ae. aegypti* colonies to investigate sub-species differences

We investigated differences among populations of the same species by examining 10 laboratory colonies of *Ae. aegypti* isolated from different geographical locations (Noah H. Rose et al. 2020), representing two subspecies: the native African subspecies *Aedes aegypti formosus* (*Aaf*) and the globally invasive subspecies *Aedes aegypti aegypti* (*Aaa*), which specializes in biting humans and breeding in human habitats (Fig.2B). The human-adapted *Aaa* subspecies diverged from its African ancestor around 5,000 years ago (Noah H Rose et al. 2023) and spread out of Africa during the Atlantic slave trade (Powell and Tabachnick 2013). While genetic signatures of *Aaa* adaptation to human habitats are emerging (Lozada-Chávez et al. 2025), observed behavioral differences between the two subspecies remain largely restricted to host preference (McBride 2016). This panel of colonies provides an ideal testbed to quantify potentially subtle behavioral differences between *Ae. aegypti* subspecies under consistent conditions. We monitored 40 mosquitoes from each colony over 10 days and performed multiscale analyses of flight behavior. All colonies exhibited the characteristic “*Aedes aegypti*” pattern with subtle, colony-specific variations (Fig.2D). To quantify differences at colony and subspecies levels, we developed two complementary statistical approaches: a local “interval-based” method using generalized mixed models and a global “pattern-based” method employing dimensionality reduction.

#### Interval-based approach reveals that *Aaa* are more active than *Aaf* during midday

For the local interval-based approach, we divided the 24-hour day into one-hour intervals and applied a generalized linear mixed model (GLMM) to each interval to analyze the average fraction of mosquitoes flying. In these models, subspecies was a fixed effect, replicate cage was a random effect, and individual days were treated as different realizations of the same variable (see methods). For each test, we report the z-score and *p*-value comparing *Aaa* and *Aaf* based on the time of day (Fig.2C). *Aaa* flew significantly more during midday (p-value < 0.01) and somewhat less during the night (p-value < 0.05), with no difference observed during morning or evening activity peaks.

We repeated this analysis for additional behaviors like sugar feeding, flight duration, and speed as described above. We developed “barcode plots” (generated automatically within the GUI app) that summarize all behavioral statistics stratified by hour, showing significant z-scores (p-value < 0.05) across all time intervals and variables (Fig. 2E). *Aaa* colonies exhibited increased sugar feeding activity, flight speed, and flight duration compared to *Aaf*. Despite significant changes at certain times, the actual differences were modest except for midday (ZT2-ZT10)flying fractions and flight duration, where differences were clearly larger than colony-level variation (Fig.S5). We further examined these two variables averaging over the same interval (ZT2-ZT10) for each colony (Fig.2F) and found a moderate positive correlation (*ρ* = 0.51), linking longer flights to increased flying percentages.

To explore differences within subspecies, we compared specific *Ae. aegypti* colonies using GLMM (KUM and RAB for *Aaf*; KPP and PHN for *Aaa*) finding small but significant variations in flight activity. For example, the PHN colony was more active at night than KPP, while RAB was more active between ZT3-ZT9 than KUM (Fig.S6A,B). These differences may reflect local adaptation of each population to specific environmental conditions.

#### Pattern-based approach confirms difference between *Aaa* and *Aaf* colonies

In addition to our GLMM analysis, we employed dimensionality reduction to uncover broader patterns in daily flight activity. Unlike GLMM, which analyzes each hour independently, dimensionality reduction captures patterns based on relative changes throughout the day. We used principal component analysis (PCA) to transform high-dimensional flight statistics into a low-dimensional space. Briefly, each day is represented as a ‘day vector,’ capturing 96 features related to the average, minimum, maximum, and standard deviation of flight activity across 1-hour intervals. PCA embeds these day vectors in two dimensions, revealing global patterns in mosquito flight activity. This approach produced distinct clusters for *Aaa* and *Aaf* colonies (Fig.2G), indicating global differences in their daily activity rhythms. We applied linear discriminant analysis to the PCA-reduced data (Fig.2H), identifying midday activity (ZT2-ZT10) as the key differentiator between the subspecies (Fig.2I), aligning with the results of our interval-based GLMM approach (Fig.2C).

Applying the local interval-based, and global pattern-based approach to quantify and visualize the activity patterns of all (sub)species, provides a comprehensive perspective on how daily rhythms vary across species, subspecies, and colonies, highlighting the integrated nature of these patterns (see Fig.S6D).

These two methods enable us to compare behavioral variables such as flight activity, sugar feeding, and flight duration within a unified framework, refining our hypotheses about observed differences. Notably, both techniques showed that human-adapted *Aaa* populations exhibit higher midday flight activity (ZT2-ZT10), a period typically marked by low activity in *Aedes aegypti*. Comparisons involving sugar feeding and flight duration, along with daily correlations, suggest that increased midday activity may be due to higher overall metabolism rather than specific circadian rhythm regulation.

### Quantifying the impact of physiological and environmental perturbations

Next, we investigated the impact of physiological and environmental perturbations on daily rhythms. Outside the laboratory, adult mosquitoes typically experience significant and complex changes in their surrounding environment and physiological states through natural behaviors like blood-feeding, or external perturbations like light pollution.

BuzzWatch provides an ideal framework to quantify the flight behavior of mosquitoes for several days or weeks before and after a given perturbation, with the potential to reveal long-term consequences. To demonstrate this idea, we present two examples of such “perturbation experiments”: one where we monitor the impact of ingesting a blood meal, and another where we examine the impact of an increase in day length.

#### Blood feeding has a long-term impact on activity patterns

We explored how a blood meal affects the daily rhythm of female mosquitoes. We tracked the activity of blood-fed female mosquitoes and non-fed controls over 19 days (Fig.3A). Consistent with previous studies (for instance in (Lima-Camara et al. 2014)), blood-feeding caused a 2-day period of strongly reduced activity. Beyond this period, we used the interval-based GLMM approach to analyze the impact of blood feeding on flight activity over the period 3-12 days post blood meal (days 10-19) (Fig.3B). Compared to the control, blood-fed mosquitoes showed a slight decrease in activity after the morning peak (ZT1-ZT5) and an increase in activity before the evening peak (ZT10-ZT11) (Fig.3C). There were no significant changes in overall flying time, sugar feeding, flight duration, or speed, suggesting subtle rhythm changes rather than major physiological shifts.

**Figure 3.**
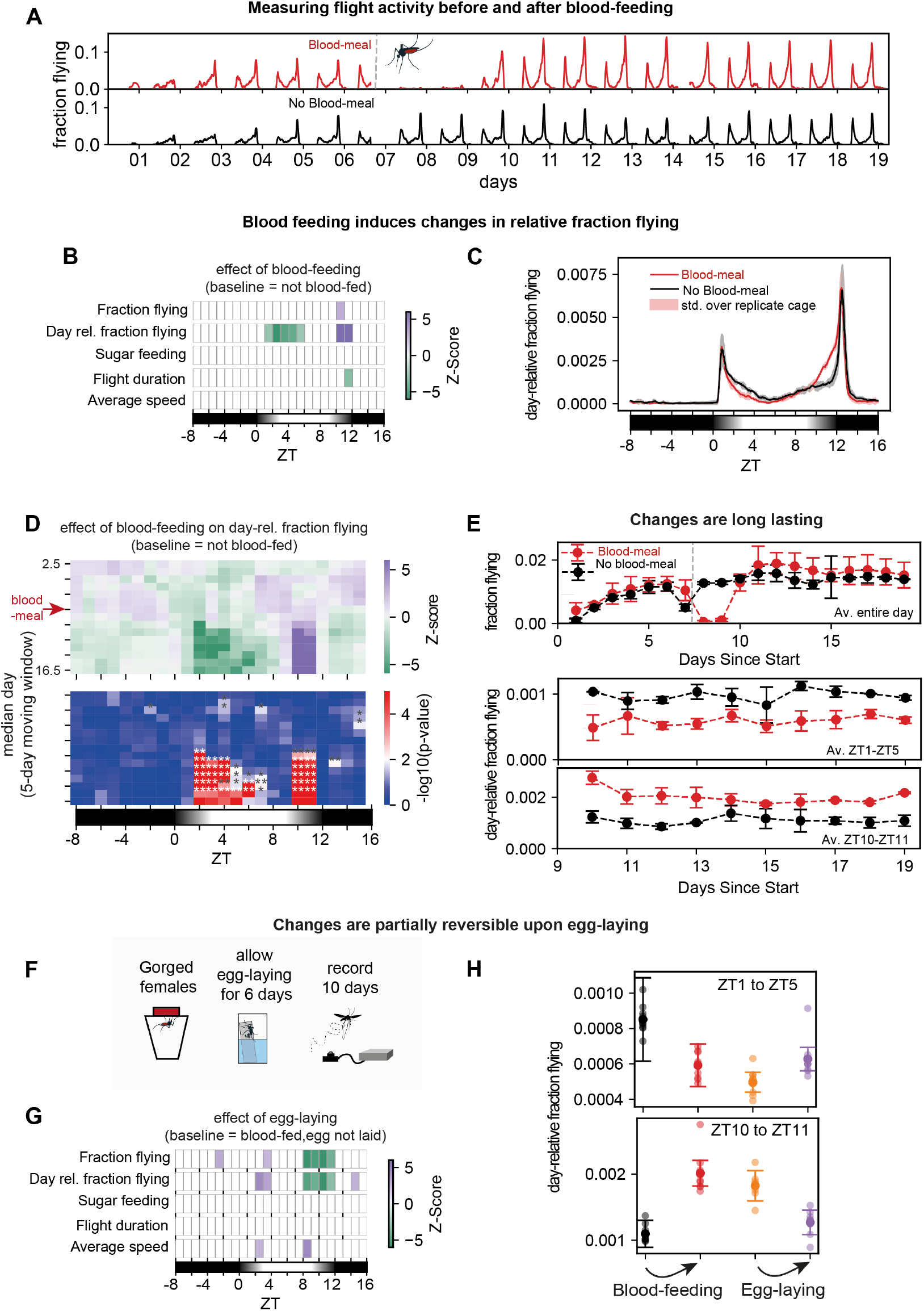
BuzzWatch allows measuring the long-term effect of physiological perturbations. **A**. Fraction of flying mosquitoes blood-fed at day 6 (red line) or not blood-fed (black line). **B**. Barcode plot of GLMM for different variables: variable names are identical to 2E. Color indicates the value of z-scores (purple for positive and. green for negative) and only z-scores corresponding to a p-value <0.05 are displayed. Data from day 10 to 19, non-blood fed control is the baseline. **C**. Day 10-19 average activity profile of blood-fed (red) and not blood-fed (black) mosquitoes. Shaded area is std. error of 2 replicate cages. **D**. Temporal heatmap for GLMM z-score (top) and p-value (bottom) on the day-relative fraction of flying mosquitoes. Columns are time of the day interval (1 hour), rows are median day interval (over a 5-day moving window). For instance, for median day 2.5 and hour ZT0, a GLMM fit was performed on data from day 1-5 with day-relative fraction flying average from ZT-1 to ZT0. In the bottom heatmap, the gradient is scaled so that p<0.01 appears in red, *p* = 0.01 in white and p>0.01 in blue. 1 star indicates 0.01<p<0.05, 2 stars p<0.01. **E**. Average fraction flying over the entire day (top), day-relative fraction flying between ZT1-ZT5 (middle) and ZT10-ZT11 (bottom) for each day, blood-fed (red) and control (black) condition. Dots are average over 2 replicate cages and error bars indicate std over these two cages. **F**. Schematic of egg-laying experiment, see Fig.S7 and methods for detailed protocol. **G**. Barcode interval based GLMM plot, baseline is “no egg laying” condition. **H**. Scatter plot comparing the average day-relative fraction flying between ZT1-ZT5 (top) and ZT10-ZT11 (bottom) over 10 days for blood-feeding and egg-laying experiments. Filled dots are average over 10 days, transparent dots are individual days, error bar indicates standard deviation over 10 days. Black is control not blood-fed, red is blood-fed – eggs not laid (from blood-feeding exp); Orange is blood-fed and eggs not laid (from egg-laying exp.), purple is blood-fed and eggs laid (from egg-laying exp.)

To assess the stability of these changes, we expanded our statistical approach, using a 5-day moving window instead of a fixed 10-day period. Each row in the heatmap shown in Figure 3D represents the median day of the window, with columns showing hourly intervals. This analysis confirms that the observed behavioral changes were stable over the 10 days after blood ingestion, with no decrease or increase in the magnitude of the change over time (Fig.3E). These findings indicate that, after a brief period of rest, blood ingestion has a moderate but persistent effect on mosquito flight activity patterns.

Next, we examined whether egg-laying could reverse these stable changes in flight activity patterns. Without an oviposition site, as was the case in the previous experiment, females may retain eggs which could explain the subtle yet long-lasting effect on mosquito behavior. To test that hypothesis, we allowed blood-fed females to lay eggs over a period of 6 days, measured their activity for the next 10 days and compared this to blood-fed females that did not have access to an oviposition site (Fig.3F, and Fig.S7A). We observed that egg-laying altered flight patterns in subsequent days, with increased activity from ZT2-ZT4 and decreased activity from ZT8-ZT12 (Fig.3G). Remarkably, this pattern mirrors changes seen after blood ingestion. Comparing the effects of blood feeding and egg-laying over the same time intervals, shows that evening flight changes were reversed, while morning changes were only partially reversed (Fig.3H). This suggests that egg-laying after blood feeding partially returns the mosquito to its original pre-blood-feeding activity pattern.

#### Complex response to photoperiod change

By feeding on humans, species like *Ae. aegypti* can be exposed to artificial light that may disrupt the natural rhythmic patterns shaped by sunlight which evolved over millions of years (Taylor and M. D. R. Jones 1969; Baik et al. 2020; Rund et al. 2020). An unanswered question is the extent of their phenotypic plasticity, specifically how quickly they adapt to these changes. As a proof of concept, we used BuzzWatch to simulate a shift from a “natural” 12-hour light/12-hour dark cycle, typical of rural areas near the equator, to a 20-hour light/4-hour dark cycle, simulating the photoperiod of urban areas. We monitored mosquitoes for 12 days under the natural cycle, then switched to the urban cycle for 10 more days (Fig.4A). After the change, mosquitoes changed from a two-peak activity rhythm (4B) to a pattern with three or four peaks (4C), illustrating their phenotypic plasticity. Among the three peaks, two clearly matched the start (ZT0) and end of the light period (ZT20), whereas an additional peak was observed around ZT14 (ZT10 in previous 12L-12D regime), not corresponding to any of the former or new light-dark transitions (Fig.4C; Fig.4D second row). Using GLMM analysis, we found that increased daylight significantly altered flight activity and reduced flight speed and duration, yet did not affect sugar feeding (Fig.4D).

**Figure 4.**
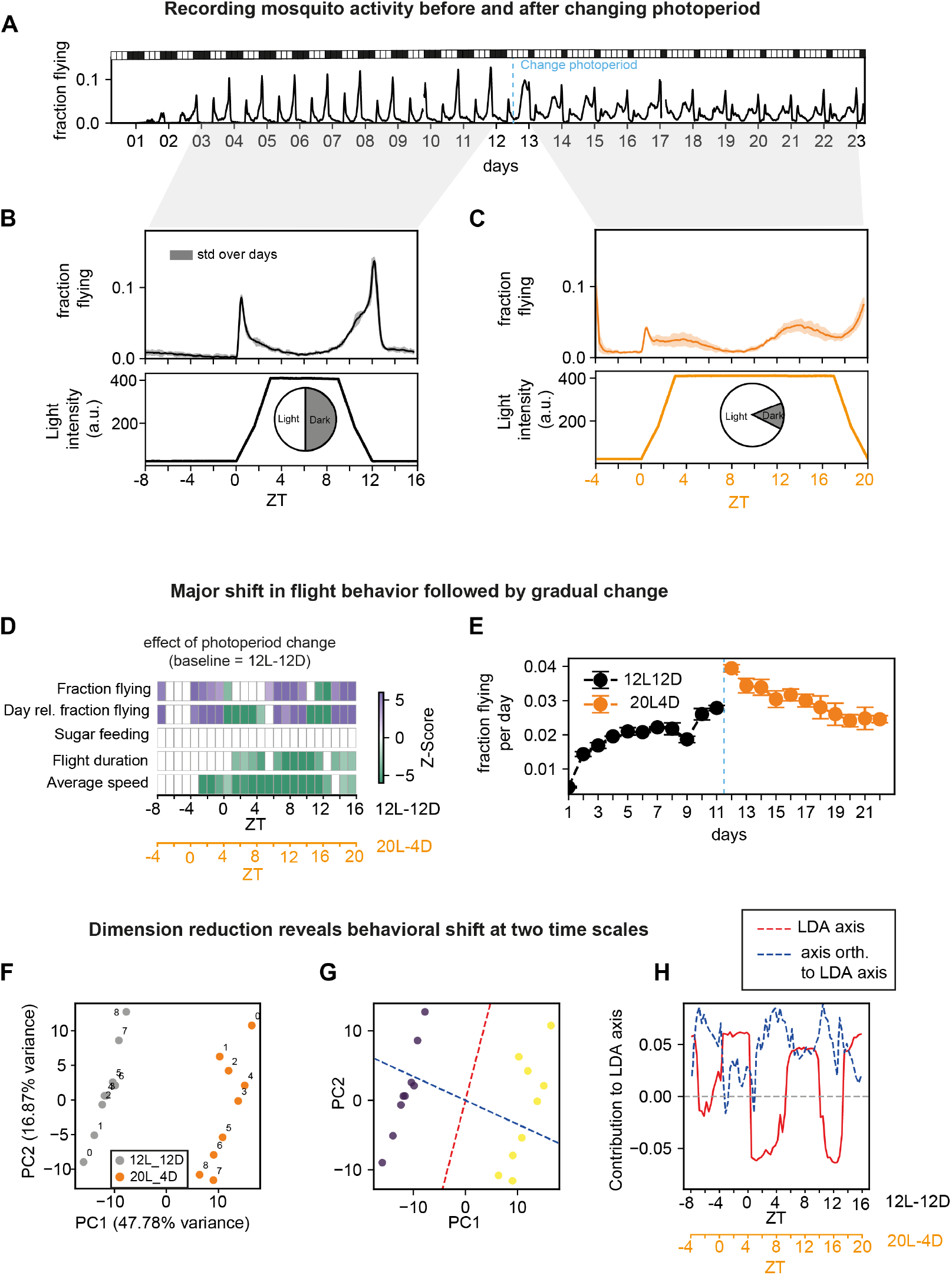
BuzzWatch reveals long-term effects of environmental perturbations. **A**. Fraction of flying mosquitoes of a representative experiment, 20 minutes moving average (black line). **B**. Top, multi-day average, in 12L12D light cycle regime (from day 3 to 12, black line). Shaded area represents std over days. Bottom, visible light intensity (blue line). At ZT0, light intensity gradually increases until ZT3, at ZT9 light intensity gradually decreases until turning off at ZT12. **C**. Top, multi-day average, in 20L4D light cycle regime (from day 13 to 23, orange line). Shaded area is std over days. Bottom, visible light intensity (blue line). At ZT0, light intensity gradually increases until ZT3, at ZT17 light intensity gradually decreases util turning off at ZT20. **D**. Barcode plot (variable same as in Fig.2E) showing changes in response to shifting the photoperiod from 12L-12D (baseline) to 20L4D. Nine days were analyzed for each condition. Only z-score for p-value<0.05 are shown. **E**. Average fraction of mosquitoes flying over a day, averaged between 2 replicate cages, error bars are std. deviation between 2 replicate cages. Black (rsp. orange) dots indicate days in 12L12D (rsp. 20L4D) photoperiod.**F**. PCA dimension reduction on day vectors, grey are days in 12L-12D and orange are days in 20L-4D. Day numbers are indicated next to each dot. **G**. Linear dispersion analysis on PCA-reduced coordinate on two groups 12L-12D (purple) and 20L-4D (yellow). Red dashed line indicates the direction of the LDA axis in PCA coordinates, blue dashed line indicates the direction of the axis orthogonal to LDA axis. **H**. Relative contributions of LDA axis (red dashed line) or orthogonal axis (blue dashed line) to hour intervals.

Beyond these immediate changes, we investigated finer behavioral shifts on longer time scales. Initially, total flight activity surged after the photoperiodic change but gradually returned to levels similar to the previous regime within 6-7 days (Fig.4E). This suggests a gradual, week-long adjustment during which mosquitoes redistribute their flight activity. To explore this further, we applied the global dimensionality reduction analysis by comparing 9 days in each photoperiod (Fig.4F). We observed a clear separation along a first axis (close to component one), corresponding to major changes in flight rhythm already detected by GLMM approach (Fig.4G-H red line). Surprisingly, in both photoperiod regimes, days structured themselves along a second axis (close to component two). Over time, days gradually move upwards along this second axis in the 12L-12D regime and downwards in 20L-4D regime (Fig.4F). The second axis (blue line in Fig.4H) primarily reflects changes in total activity (as seen in Fig.4E), alongside subtle adjustments in flight rhythm. A complete absence of rhythm change would appear as a flat blue line in Fig.4H. A possible interpretation is that mosquitoes initially aligned with the 12L-12D cycle retain a “memory” of it, before gradually shifting under new conditions.

Overall, this study revealed two aspects of the plastic response of mosquitoes to photoperiodic change: rapid rhythm changes within 1-2 days, and slower, refined adjustments over weeks. These results underscore the complex temporal aspect of phenotypic plasticity of mosquito flight behavior in response to environmental changes.

### Measuring the response to pulse of host-cues at different times of the day

Next, we used our platform to quantify mosquito responses to short pulses of host cues. Previously, we measured “spontaneous” flight activity without host cues. While it is unclear if this is driven by a quest for hosts, increased activity may certainly contribute to host-seeking by raising the chances of encountering a host or detecting its cues. However, when mosquitoes are inactive, such as at night or midday (Fig.1D), their response to host cues is uncertain. Here, we focused on CO_2_ and heat for their consistency and reproducibility (Sorrells et al. 2022; McMeniman et al. 2014). We designed a system controlled by a Raspberry Pi board to deliver controlled pulses of CO_2_ and heat at user-defined intervals. This system includes a CO_2_ canister, a “gas mixing” chamber with a solenoid valve, an air pump, and a Peltier module to heat a small area of the cage (Fig.5A). We observed that 30 seconds of CO_2_ followed by 4.5 minutes of heat effectively attracted mosquitoes for several minutes. While pulse parameters can be adjusted, we used this 5-minute sequence consistently and refer to it as a “single host-cue pulse.”

**Figure 5.**
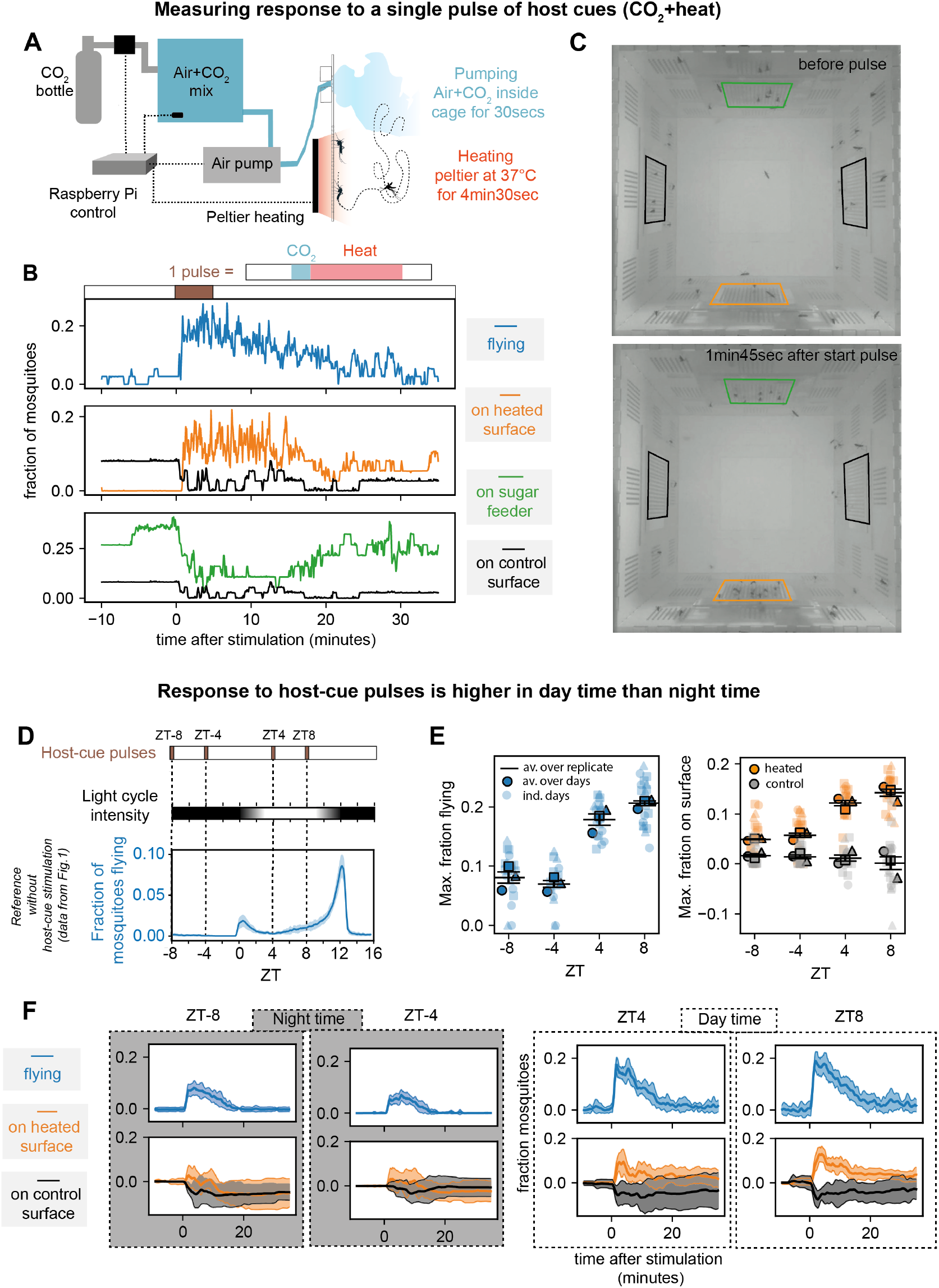
BuzzWatch allows investigating response to pulses of host-cues at different times of the day. **A**. Delivery of host-cue pulses is controlled by a raspberry pi, with the following sequence : (1) opening of the solenoid valve for 5 minutes, CO_2_ goes from the CO_2_ canister to a mixing box (increasing the concentration from 400ppm to 1000ppm) (2) Mix of Air+CO_2_ is pumped for 30sec from the mix box to the cage containing mosquitoes, at approximately 2.5 liter/minute (equivalent to 1/3 of the cage volume). (3) Peltier placed 5mm behind the surface of the cage is heated at 37°C for 4min30sec. **B**. Response to a single pulse of host-cues (CO_2_ and heat) starting at t = 0min and finishing at t = 5min (brown bar), for a cage of 40 unfed *Ae. aegypti* females (8-10 days old at the start of the pulse and kept in the cage with 12L12D for 5 days). Moving average with 3-second window of the fraction of mosquitoes either flying (blue line); on the heated surface (orange line, from orange region of interest (roi) in 5D); on the sugar feeder (green line, from green roi in Fig.5D) and control surface (black line, from black roi in Fig. 5D). Black lines in the middle and bottom graph are the same (control for reference) **C**. Snapshot images of movies, before the initiation of a host-cue pulse sequence (top, t =-2min) and during the pulse (bottom, t =+1min45sec). Colored rectangle overlayed indicates the roi used to track the number of mosquitos on the heated surface (orange); the sugar feeder (green); control surfaces (black). **D**. Timing of host-cue pulse stimulation. (brown bars, CO_2_+heat), 4 times per day over 9 consecutive days, at night times (ZT-8, ZT-4) and at day times (ZT4, ZT8). Three replicate cages of 40 *Ae. aegypti* females. Bottom, fraction flying averaged over 32 days from Fig.1, *Ae. aegypti* not stimulated by host-cue pulses. **E**. For each host-cue pulse, fraction of flying and fraction on the heated surface (or control) was averaged with 10sec rolling window. Then the max between t=0 (the beginning of the pulse) and t = 10min (5min after the end of the pulse) was computed for the fraction flying (blue), the fraction on the heated surface (orange) and the fraction on the control surface (black). The transparent dots are max value of a single pulse, different shapes represent different replicate cages; plain dots with black edges are average of all the pulses at a given time over 9 days for a given replicate. Horizontal black line is the average over the 3 replicate cages for a given pulse time, error bars are s.e.m. One-sided t-test on averages per replicate (n=3) for flight activity (left) ZT-8 - ZT-4 p-value = 0.439; ZT-8 - ZT4 p-value = 0.010; ZT-8 - ZT4 p-value = 0.0006; ZT4 - ZT4 p-value = 0.074. For difference between heated (orange) and control surface (grey) (right) : ZT-8 : p-value = 0.062; ZT-4 : p-value = 0.019; ZT4 : p-value = 0.0017; ZT4 : p-value = 0.0002; For difference between heated surface at different times (orange) (right) : ZT-8-ZT-4 p-value = 0.029; ZT-8-ZT4 p-value = 0.002; ZT-8-ZT4 p-value = 0.002; ZT4-ZT4 p-value = 0.184. **F**. Response to host pulse for a single replicate cage, curves are averaged over 9 different pulses of 9 different days. Shaded is standard deviation over the 9 pulses.

#### Response to a single pulse of host cues

Starting with a “single pulse” experiment, we examined mosquito responses to CO_2_ and heat. We quantified the fraction of flying mosquitoes and their distribution over several surfaces (heated, sugar-feeder, and controls, see Fig.5B-C). Immediately after the pulse, flight activity surged, slowly decreasing over several minutes (Fig.5B top). Mosquitoes quickly gathered at the heated area, while other areas showed no change, or a reduction in occupancy (Fig.5B middle). While not encompassing the full complexity of blood feeding behavior, this experiment shows that monitoring mosquito distribution over time reliably quantifies responses to host cues serving as a proxy for host-seeking behavior (Supp. Movie 1, mosquito inserting their proboscis through the acrylic holes in response to the heated surface).

#### Host-seeking response at different times of the day

We asked how the response to pulses of host cues varied throughout the day. As previously observed, in the absence of host cues *Ae. aegypti* flight activity is virtually absent during midday and nighttime (Fig.2, Fig.5D). To explore if *Ae. aegypti* responds to host-associated cues during these apparent “resting” times, we tested four stimulations daily at ZT-8, ZT-4, ZT4 and ZT8 over nine days to quantify the behavioral response (Fig.5D). We stimulated triplicate cages containing 40 *Ae. aegypti* females with this pulse regime, measuring total flight activity and attraction to the heated surface (as in Fig.5B). We analyzed peak responses in a 10-minute window after the pulse, smoothing data with a 1-minute average (Fig.5F). Responses were consistent for a given time of day, differed greatly between day and night, and varied minimally within those periods (Fig.5E-F). Flight activity and attraction were 2-3 times higher during the day (Fig.5E). Importantly, even at night 10% of mosquitoes showed increased activity in response to the stimulus, a level of activity comparable to the evening peak without stimuli. However, nighttime attraction to heat was modest, sometimes similar to control surface fluctuations, unlike daytime responses.

By expanding BuzzWatch with a module to consistently deliver host-associated cues, we showed that strength of the responses to CO_2_ and heat varies throughout the 24-hour cycle, yet responses are consistent across days at specific times. Even at times when mosquitoes typically rest, they respond measurably to stimulation, although *Ae. aegypti’s* host-seeking response is approximately twice as high during the day compared to night. This suggests that the host-seeking response is heavily influenced by circadian rhythms, in ways that cannot be solely explained by baseline flight activity. This proof of concept demonstrates that BuzzWatch can effectively characterize mosquito responses to host cues.

## DISCUSSION

BuzzWatch establishes a novel approach to longitudinally study multi-scale temporal aspects of mosquito flight activity. We developed a comprehensive methodological pipeline that encompasses designing cages, setting up mosquito recordings, automatically extracting flight tracks from video footage, and analyzing these tracks to effectively quantify mosquito activity over weeks in a tightly controlled, user-defined environmental setting. BuzzWatch enables researchers to gather longitudinal, rich population-level datasets, allowing comprehensive analyses of mosquito daily rhythm, phenotypic plasticity, and responses to environmental cues.

Flight is fundamental to initiating key behaviors in adult mosquitoes, such as sugar feeding, host seeking, mating, and oviposition. Monitoring such behaviors over weeks presents the challenge of tracking flight effectively. Previous studies used high-precision multi-camera 3D flight tracking to characterize short-term mosquito flight behaviors such as attraction to human cues during host-seeking and environmental interactions (Gupta et al. 2024) (van Breugel et al. 2015). However, sophisticated multi-camera setups are not used to characterize behaviors over longer periods of time as it would quickly generate impractical amounts of data requiring massive computational resources for analysis. The complexity of multi-camera 3D tracking setups furthermore requires specific laboratory expertise and considerable financial resources. While obtaining high-resolution tracking data of the intricate 3D maneuvers characteristic of mosquito flight is essential in some contexts, a wide variety of behaviors can be assessed without this level of detail. With BuzzWatch, we therefore opted for the approach of using a single low-cost camera to monitor flight behaviors of cohorts of mosquitoes, and developed analysis tools to extract key behavioral metrics from movies spanning hundreds of hours. While the full 3D flight track is reduced to a 2D projection in our setup, it allows accurate long-term assessment of activity patterns, sugar feeding behavior, responses to host cues, and certain flight statistics. The BuzzWatch setup is low-cost and can be constructed from readily available materials, making it a scalable approach to run several month-long replicate experiments in parallel, a capacity that is out of reach for current multi-camera 3D flight tracking setups.

In contrast to camera-based approaches that monitor freely flying mosquitoes, tube-based locomotor activity monitoring systems like the commonly used Drosophila Activity Monitor (DAM) (Chiu et al. 2010) use motion induced beam breaks to quantify activity. These systems, and camera-based derivatives (e.g. (Sondhi et al. 2022) and Araujo, Guo, and Rosbash 2020), are easy to set up and compatible with monitoring mosquitoes over several days. However, activity in these systems is only detected when a moving insect blocks a light beam (likely underestimating true activity, see (Keleş et al. 2025)). The obtained data is therefore limited to counting “active” versus “inactive” states while not providing locomotion statistics (e.g. duration and speed of movement), nor do they provide insight regarding when an insect visits a specific area of interest such as a sugar feeder, or a surface presenting host-associated cues. These assays furthermore typically constrain animals in narrow tubes or 24-well plates, severely restricting movement and preventing flight, fundamental to many mosquito behaviors. Compared to multi-camera 3D flight tracking approaches on the one hand, and beam-block activity monitors on the other, BuzzWatch provides an affordable and scalable alternative for accurate long-term activity monitoring of cohorts of freely flying mosquitoes, and assessment of key behaviors such as responses to host cues, and sugar feeding. BuzzWatch incorporates a comprehensive downstream pipeline to rigorously analyze mosquito behavior across multiple timescales. To empower researchers to rigorously test hypotheses using advanced statistical tools, we provide an extensive GUI application that facilitates statistical analysis, enabling reproducible comparisons between different conditions and investigating emerging patterns in temporal data. We implemented two complementary approaches: a local “interval-based” analysis using GLMMs and a global “pattern-based” analysis using dimensionality reduction techniques. With BuzzWatch we establish an integrated open-source hardware and software pipeline from cage design to statistical analysis of behavior. While we optimized each step for our studies, we adopted a modular and flexible approach using generic and low-cost parts to enable adoption and adaptation by others.

### Origins and implication of changes in mosquito flight-activity patterns

By comparing diverse populations of *Ae. aegypti*, we found that the *Aaa* subspecies exhibits more midday flight activity and increased sugar feeding compared to the *Aaf* subspecies. All experimental colonies had been maintained in laboratory conditions for only 15-30 generations (see Table1 and supplementary file 1). Previous studies on rhythmic activities used older lab colonies of *Aaa*, complicating direct comparisons (Ajayi et al. 2024). Known differences between *Ae. aegypti* subspecies include higher susceptibility to arbovirus infection for *Aaa* (Aubry et al. 2020) and increased preference for human blood meals and human-associated oviposition sites (Powell and Tabachnick 2013; McBride 2016). Our data suggest that higher midday flight activity of *Aaa* reflects longer flights and increased sugar feeding, indicating possible physiological differences. Whether this behavior is directly related to the adaptation of *Aaa* to human habitats remains to be investigated. To gain more insight into the origin of the changes in flight rhythm, future research could leverage genomic data comparing multiple *Aaa* and *Aaf* populations (Lozada-Chávez et al. 2025) to identify genes associated with flight rhythm or metabolic differences. It would be interesting to investigate whether the higher midday flight activity of *Aaa* translates into elevated response to host-cues during midday. Indeed, recent studies show that temporal patterns of locomotor activity and host-seeking response are determined by genetic factors in *Ae. aegypti* (Dong et al. 2024).

### BuzzWatch and phenotypic plasticity in mosquitoes

In response to physiological changes, like blood ingestion, and environmental shifts, such as a sudden photoperiod increase, we observed a rapid initial shift in behavior followed by subtle long-term adjustments or stability. Further research is needed to determine if these findings reflect the general robustness and plasticity of mosquito behavior in changing environments. Specifically, BuzzWatch is a potent tool for addressing public health issues linked to mosquito phenotypic plasticity. Indeed, as major vectors of human pathogens, *Ae. aegypti* and *Anopheles* mosquitoes are often exposed to control measures like insecticides and traps. Several field studies have reported behavioral shifts that enable mosquitoes to escape these control measures, threatening their long-term efficacy (Carrasco et al. 2019). To understand how such “behavioral resistance” to interventions evolves, it is crucial to determine whether these traits are constitutive or induced (i.e., phenotypically plastic). We foresee BuzzWatch as a promising tool to rigorously quantify behavioral resistance to interventions like insecticides and shedding light on its origins.

*Ae. aegypti* and *Anopheles* mosquitoes also face pathogen infections over approximately two weeks, during which pathogens replicate and spread through the organism. The long-term effects of these infections, especially regarding arboviruses (Maire, Lambrechts, and Hol 2024), remain unclear. The BuzzWatch platform could be instrumental in tracking the flight behavior of *Plasmodium*- and arbovirus-infected mosquitoes in the weeks following an infectious blood meal.

Overall, BuzzWatch represents a significant advancement in the study of mosquito behavior, providing an integrated, open-source platform that enhances our understanding of phenotypic plasticity and adaptive responses and holds promise for contributing to effective public health strategies in combating mosquito-borne diseases.

## METHODS

### Cage design and fabrication

Cages were created from acrylic (Richardson) using a laser cutter (Trotec Speedy 300). Cage designs were adapted from boxes created using boxes.py (https://boxes.hackerspace-bamberg.de/). Cage parts and supports were either glued with commercial superglue or with dichloromethane-based acrylic adhesive (Acrodis, Sun-clear ref029019). Three millimeter thick acrylic was used for all cage parts, except for the perforated side windows covering the sugar feeders which were made out of 1.5 mm thick acrylic in order to allow mosquito to easily reach the sugar cotton by inserting their proboscis through 1 mm diameter holes. For the raspberry pi camera to record mosquito activity, transparent acrylic has to be used for the bottom part of the cage. For all the other parts, either transparent or white diffusive acrylic can be used to improve image quality. In order to create a uniform visual background, white paper was placed on top of the cage. An infrared LED light source (Andoer IR49S LED, usb-c powered) was placed 20 cm above each cage, to ensure constant and homogenous lighting for video recording independent of variable visible light intensities. A Raspberry PI V2 NOIR 8MP camera was placed 20 cm below each cage to record the activity of all mosquitoes within the cage. Camera focus was adjusted manually so that the both the bottom and the top of the cage were equally sharp, to minimize bias in tracking mosquitoes located at different distances from the camera. A 850 nm infrared long-pass filter (Thorlabs FEL0850) was placed in front of the camera lens so that the camera sensor only received infrared light, blocking all visible wavelengths. This assures that background image intensity and contrast remain constant throughout the days, facilitating analysis. See the BuzzWatch website for design files and extensive tutorial on construction. Four cages were placed in an enclosure (600mm*750mm*750mm) constructed from 3 mm white opaque acrylic and aluminum rail profiles (see Fig.S1). Two RBG LEDs strips connected to a Raspberry Pi (NeoPixel RGB 1m 60 LEDs ADA1138) were installed inside the container to control the daylight cycle. A python script using the *neopixels* module controlled the light intensity of all LEDs in the strip, according to a pre-defined day/night cycle with 3-hour gradual dimming. Temperature, Humidity sensors (DHT11) as well as a light sensor (TSL2591) were placed inside the container for continuous logging.

### Video recording and storage

Each camera was connected to a Raspberry Pi 4B+ micro-computer, which was programmed to automatically record videos in h264 format. Videos were stored in 20-minute segments on the micro-SD card. Cron scheduling rebooted the Raspberry Pi automatically every 12 hours, converted the h264 files on the SD card to mp4 format, and transferred all mp4 files to an external SSD. Video recording settings were manually adjusted before the start of recording using the Python *pirecorder* module (Jolles 2020) interface and kept constant during the experiment. A Python script utilizing the *pirecorder* module handled the recording and conversion to .mp4. At the conclusion of the experiment, all .mp4 videos (approximately 200-400 GB) were transferred to a server or a local workstation for analysis. An example image of the SD-card with all settings is available for download.

### BuzzWatch GUI app for mosquito tracking and data analysis

We developed a cross-platform graphical user interface app in Python to streamline both the tracking of mosquitoes and the subsequent analysis of mosquito behavior.

### Mosquito tracking

Our analysis pipeline employs standard computer vision techniques, such as background subtraction and thresholding, to segment mosquitoes from the cage background. A clean background image is generated by averaging 100 frames from 100 consecutive 20-minute videos, spanning over 30 hours, ensuring no mosquitoes are visible in the background image. This clean background is crucial as mosquitoes typically move at least once within 24 hours. Two distinct parallel operations are performed to track resting and flying mosquitoes. Resting mosquitoes are identified by subtracting the background from all frames, followed by blob detection to segment and track centroids. A greedy algorithm processes these centroids to form tracks of resting mosquitoes, capturing both stationary and slow-moving individuals. For flying mosquitoes, the previous frame is subtracted—a standard motion detection method—followed by segmentation and centroid tracking optimized for flying mosquitoes. A custom algorithm links tracks of resting and flying mosquitoes to identify take-off and landing events. The algorithm performs a local search to find matching pairs, assuming spatial proximity between the end of a resting track and the start of a flight track for take-off events, and vice versa for landings. This bootstrapping procedure is iterated five times to enhance matching accuracy, an minimize identity swaps. Importantly, identity swaps do not compromise the extraction of population-level variables, such as the fraction of mosquitoes flying or resting on specific surfaces, such as sugar feeders. Final mosquito tracks are stored as pickle files for each analyzed video, containing coordinate data and IDs for 40 mosquitoes at 25 frames per second, with file sizes typically ranging from 20–30 MB.

### Data analysis, exploration, and visualization

Unless specified otherwise, for all population level data (fraction flying; fraction sugar feeding) we applied the same normalization procedure: time-series were resampled to 1-minute intervals, normalized by the total number of alive mosquitoes, and smoothed by a 20-minute rolling window. For the “day-relative fraction flying” we further normalized by the sum of the fraction flying over a given day from 0h00 to 23h59 (area under the curve).

### Interval-based GLMM analysis

To assess the impact of various conditions on mosquito activity at specific times of the day, we employed a generalized linear mixed model (GLMM). We divided each day into short time intervals (e.g., 20 minutes) and calculated the average value of mosquito activity variables within these intervals, such as the average proportion of mosquitoes flying between 12:00 and 12:20. The model is formulated as follows:

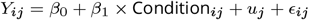

Where:

- *Y*_*ij*_ represents the outcome for the *i*-th observation within the *j*-th experiment.
- *β*_0_ is the intercept, representing the baseline level of activity.
- *β*_1_ ×Condition_*ij*_ denotes the fixed effect of the condition being tested (e.g., blood-fed vs. non-blood-fed).
- *u*_*j*_ is the random intercept for the *j*-th experiment, modeled as *u*_*j*_ ~ 𝒩 (0, *σ*_*u*_), accounting for variability between replicates.
- *ϵ*_*ij*_ is the residual error, assumed to be *ϵ*_*ij*_ ~ 𝒩 (0, *σ*^2^).

This model does not explicitly account for specific day-to-day effects; values from different days are treated as separate realizations of the same statistical variable. Each mosquito activity variable for each time interval is modeled independently, meaning cross-correlations between time intervals or effects spanning different intervals are not directly addressed.

To define and fit each Generalized Linear Mixed Model (GLMM), we used the Python module `MixedLM` from `statsmodels.regression.mixed_linear_model`, with `method` = ‘lbfgs’, `maxiter` = 500, and used the summary output to extract the coefficient, standard error, p-value, and z-score (and confidence interval). The z-score is defined as the coefficient divided by its standard error.

As an example, for Fig. 2 and the *Ae. aegypti* sub-species analysis, the GLMM is formulated as follows:

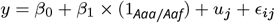

where:

- *β*_0_ is the intercept,
- *β*_1_ × (1_*Aaa/Aaf*_) represents the fixed effect,
- *u*_*j*_ is the random effect,
- *ϵ*_*ij*_ is the residual noise.

### Pattern-based dimension reduction analysis

To compare activity patterns between days, replicates, or conditions, we used dimensionality reduction. As for the GLMM model, we divided each day into non-overlapping intervals of one hour (interval can be changed in the GUI). Each day was represented as a high-dimensional vector consisting of 96 features, detailing the average, minimum, maximum, and standard deviation of flight activity variables for each 1-hour interval. We referred to these as “day-vectors”. To facilitate comparison between different day vectors, we normalized the data for mean and standard deviation (across the ensemble of day vectors considered) and applied dimensionality reduction using principal component analysis (PCA) (in the GUI app one can alternatively use UMAP or t-SNE). This allowed us to visualize day-vectors in a 2D space, where the two axes are the principal components capturing the greatest variance. Two day-vectors positioned closely in this reduced space indicate a “globally” similar daily activity pattern. Note that the relative positioning depends on the specific PCA axes that were computed. Importantly, PCA is independent of labels assigned to day-vectors (like different colonies). Therefore, to compare two groups of day-vectors, such as different subspecies, we applied Linear Discriminant Analysis (LDA) on the PCA-reduced coordinates. This identifies the axis that best separates the two groups. To interpret this separation, we back-transformed the LDA axis into time-of-day coordinates. This was possible because PCA axes are linear combinations of the original 96 dimensions. The relative contributions (weights) of each dimension were considered, and contributions were grouped by time intervals by selecting the highest contribution from the four statistics (average, min, max, and std) for each interval. This way, contributions can be represented as function of the time of day. This approach, using PCA-reduced space, offers a robust way to discern differences between groups by accounting for overall variance and structure among all day-vectors. This procedure can be applied to any PCA-derived axis, such as the first or second principal components, the LDA axis, or the axis orthogonal to the LDA axis, as illustrated in Fig.3G-H.

### Mosquito rearing

The mosquito species/strains used in this study are described in Table 1. *Ae. aegypti* and *Ae. albopictus* mosquitoes were reared at Institut Pasteur. Eggs were hatched in dechlorinated tap water in a vacuum chamber for 45 minutes. Then 200 newly hatched larvae were transferred to a plastic tray filled with 1.5 liter water. Larvae were kept at 28°C, 40% relative humidity on a standard diet of fish food (Tetramin, Tetra). Fish food was added on the first day after hatching, and a second time after 3 days. After 6-7 days, pupae were collected in a 10cm wide, 15cm high plastic container with clean water and placed in BugDorm-1 cages (BugDorm) for adult emergence. Emergence cups were removed after 4 days. Adult *Aedes* mosquitoes were maintained at 28 ± 1^°^C, 75% relative humidity, and a photoperiod of 12 hr light : 12 hr dark in 30 × 30 × 30 cm screened BugDorm-1 cages having continuous access to 10% sucrose, without separating males and females. *Anopheles stephensi* were reared at Radboudumc, Nijmegen under similar temperature, humidity, and diurnal cycle conditions, with continuous access to 5% glucose.

### BuzzWatch experiment

Three to five days after emergence, *Aedes* mosquitoes were aspirated and knocked-down on ice for 10 minutes. Females were selected and counted in a petri dish on ice. Once all females were sorted in one petri dish, they were transferred inside an acrylic BuzzWatch cage that was immediately closed with screws. *An. stephensi* females were aspirated directly into the BuzzWatch cage. The sugar feeders were filled with cotton soaked with 10% sucrose solution, excess liquid was absorbed with paper and feeders were placed carefully on the side of each cage (Fig.S3), supported by elastic bands. Cages were placed inside the setup (Fig.S1) for recording.

#### Specific protocol Fig.1 and Fig.2

One cage containing 40 *Ae. aegypti* females strain KPP aged 3-5 days old was monitored for 32 days. A single sugar feeder was placed on the side of the cage and was changed weekly. One cage containing 40 Ae. aegypti females strain KPP aged 3-5 days old was monitored for 32 days. A single sugar feeder was placed on the side of the cage and was changed weekly.

#### Specific protocol Fig.3 blood feeding (see also Fig.S7 for egg-laying)

For the blood-feeding experiment, 4 cages each containing 40 *Ae. aegypti* KPP females aged 3-5 days old were monitored in parallel for 19 days. At day 6, all cages were removed from the recording platform and brought to the blood-feeding room for 1 hour (17h-18h). Rabbit blood was provided with a membrane feeder heated to 37C to Cage #1 and Cage #4; 39/40 (Cage#1) and 40/40 (Cage#4) mosquitoes engorged (verified by visual inspection). No blood was provided to control Cage #2 and #3. All 4 cages were returned to the BuzzWatch platform and recording resumed.

For the egg-laying experiment, mosquitoes were reared in large bug dorm cages with males and females together for 3-5 days. Females were then aspirated, kept on ice in a 4°C fridge for 10 minutes, and sorted with tweezers. Approximately 360 females were sorted into boxes (60 females per box) and maintained without sugar cotton overnight. The next day, all mosquitoes were offered a blood meal using a membrane feeder. Immediately after feeding, boxes were placed on ice for sorting. Well-fed females, distinguished by visible blood in the abdomen, were sorted and placed in small bug dorm cages (60 per cage). Cotton soaked with 10% sucrose was placed on top of each cage to allow sugar feeding while preventing egg laying on the cotton (which may occur if it is placed inside the cage). After two days, egg cups with absorbent paper (ref) were placed in two cages to allow egg-laying, while nothing was added to the control cages. After four more days, the egg cups were removed, and the cages were placed on ice in a 4°C fridge for 10 minutes. Forty females per cage were sorted, placed in a petri dish, and then transferred to a BuzzWatch cage. The sugar feeders were changed weekly week during the subsequent 21 days of continuous monitoring. Inspection of the sugar feeder cotton (placed behind a 1mm diameter and 1.5mm width acrylic pierced grid) showed no eggs, consistent with previous observations that mosquitoes do not lay eggs through these holes.

#### Specific protocol 4 light cycle change

Four cages each containing 40 *Ae. aegypti* KPP females aged 3-5 days old (from the same rearing batch, prepared as in protocol 1) were monitored in parallel for 23 days. At day 13, the light-cycle was changed from 12h light – 12 hours dark to 20 light – 4 hours dark. One sugar feeder was used per cage, changed weekly.

#### Specific protocol 5 host-seeking

A cage containing 40 *Ae. aegypti* KPP females aged 3-5 days old were prepared as in the Fig.1 protocol. After 5 days adaptation to the set-up, the cage was submitted to host-cue pulses for 5 minutes at 4 fixed times (ZT-8; ZT-4; ZT4 and ZT8) for 9 consecutive days. A sugar feeder was placed opposite to the host-cue delivery window, changed 1 day before the host-cue stimulation subsequently not replaced for 10 days. The experiment was repeated 3 times (different mosquitoes and different replicate cages).

## Acknowledgements

We thank Catherine Lallemand (Pasteur) and Geert-Jan van Gemert (Radboud UMC) for assistance with mosquito rearing. We are grateful to all members of the Lambrechts and Hol labs for their insights and feedback during this project, and comments on the manuscript. We are grateful to Noah Rose, Carolyn McBride, Jewelna Akorli, Sampson Otoo, Joel Lutomiah, Rosemary Sang, Massamba Sylla, Martin Mayanja, Julius Lutwama, Alain Kohl, Alongkot Ponlawat, Veasna Duong, Claudia Romero-Vivas, Isabelle Dusfour, and Anubis Vega-Rúa, for their contributions in establishing the *Ae. aegypti* colonies. We thank members of the FabLab of Institut Pasteur for technical assistance with laser cutting acrylic and structural elements of the container.

## Contributions

T.M : Conceptualization, Resources, Data curation, Software, Formal analysis, Validation, Investigation, Visualization, Methodology, Writing - original draft.

Z.W. : Data curation, Formal analysis, Validation, Methodology (Performed experiments and analysis with *Anopheles* mosquitoes and helped with development of the method)

L.L. : Resources, Project administration, Writing - review and editing, Funding acquisition.

F.J.H. : Conceptualization, Investigation, Resources, Writing - original draft, Project administration, Funding acquisition.

## Code and Data availability

Webpage with tutorials: https://theomaire.github.io/buzzwatch/

Graphical user app repository: https://github.com/theomaire/BuzzWatch_app_v2

Repository with all processed data to run analysis in the BuzzWatch analysis app :

## Supplementary Figures

**Figure S1.**
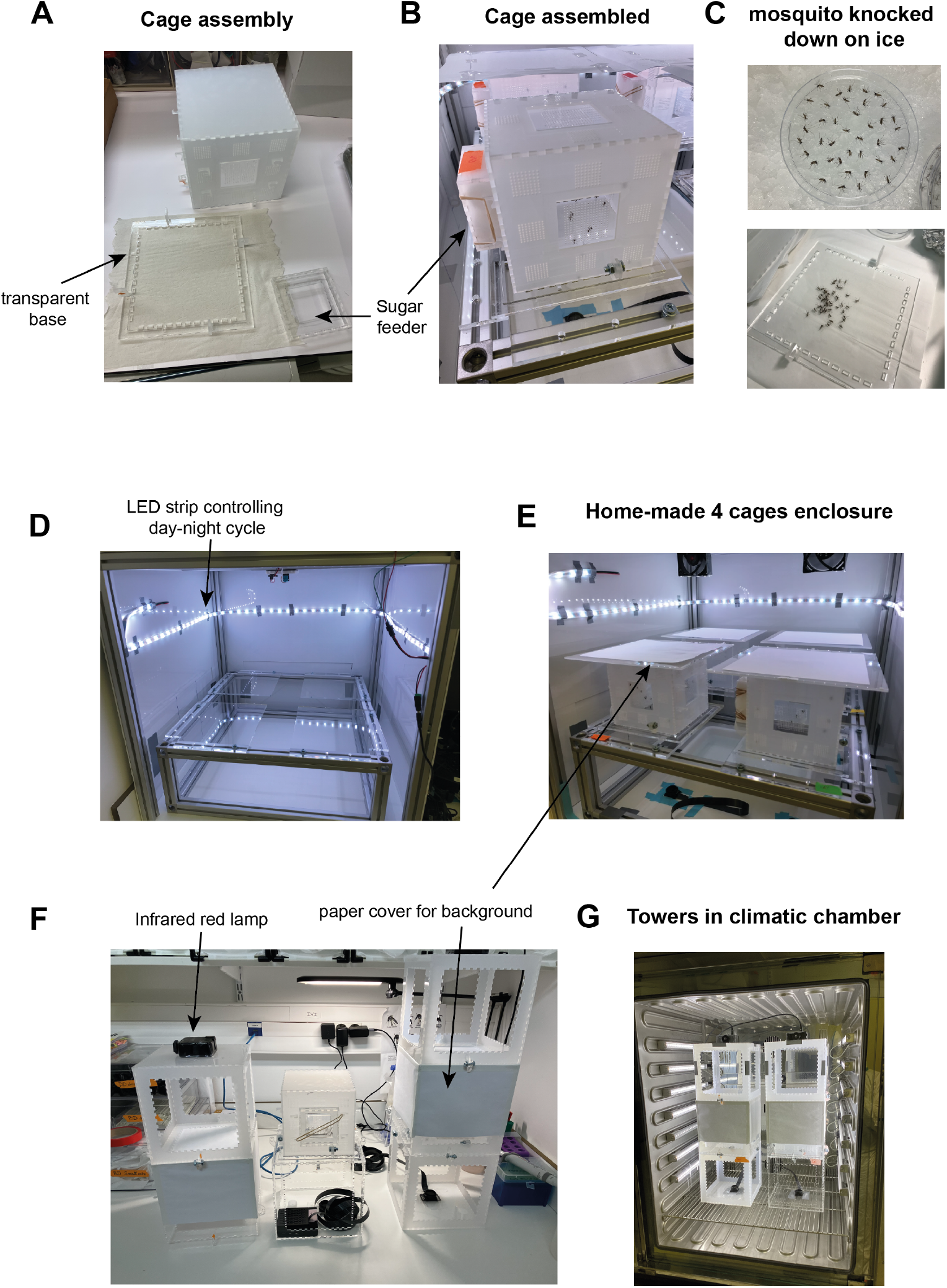
Pictures of different versions of the BuzzWatch setup. **A**. Unmounted cage, base, and sugar feeder. The cage has a transparent base so that all mosquitoes are visible to the camera placed beneath the cage. The base can be easily unmounted and attached with screws. **B**. Cage with mosquitoes and sugar feeder attached. The cage is made of 3mm acrylic except for the square “windows” on each side. These windows are 1.5mm or 2mm wide and perforated with a grid of 1mm diameter holes to allow mosquitoes to insert their proboscis for sugar feeding. Cotton soaked in a 10% sucrose solution is placed on the other side of the window. Filling the sugar feeder with soaked cotton and fixing it to the windows with elastic bands allows mosquitoes to feed ad libitum. After one week, the cotton is usually still wet, but it is preferable to change it to avoid the accumulation of fungi and mold. The rest of the cage is also perforated with 1mm diameter holes to allow air circulation. This version was used for experiments at Institut Pasteur (all data from Fig. 1-5, except for the experiment with Anopheles conducted at Radboud UMC). **C**. C **D**. Enclosure without cages and camera installed. A white LED strip creates the artificial day/night cycle, and sensors for temperature, humidity, and light are attached near the top of the enclosure. LEDs and sensors are connected to a Raspberry Pi, controlling LED intensity and logging environmental data. Aluminum beams and white opaque acrylic panels isolate four cages from the room environment, maintaining a uniform light, humidity, and temperature environment. Silent 12V DC fans run continuously on top of the container to mix the air and maintain a uniform temperature. **E**. The enclosure can accommodate four mosquito cages. In environments without external humidity control, a tray filled with tap water can be placed beneath the cages to maintain high humidity. While closed with infrared light on, the humidity remains between 60-70% over weeks, and the temperature between 24-26°C, with little fluctuation between day and night due to the white LED strip heating (see supplementary material). Acrylic lids covered with white paper were placed on top of each cage to improve the uniformity of the background image during video recording (white paper produces a uniform background that contrasts well with mosquito bodies). **F**. On the left, the unmounted tower; on the right, the complete tower with three levels: the base part holds the camera at the bottom and supports the mosquito cage; the middle part is the mosquito cage surrounded by white paper to make the background image more suitable for image analysis (mosquito detection is facilitated if the background is white and uniform). The top part holds the infrared light. The three parts can be removed or attached by screws on the sides. Once attached, the camera view is fixed relative to the mosquito cage and infrared light, so the background image is little affected by displacement of the entire tower. This is an individual tower version, with complete construction details available at https://theomaire.github.io/buzzwatch/construct.html. **G**. Two fully mounted and functional BuzzWatch towers are placed in a climatic chamber. Light, temperature, and humidity are controlled by the climatic chamber. Raspberry Pi units recording videos are placed outside the climatic chamber.

**Figure S2.**
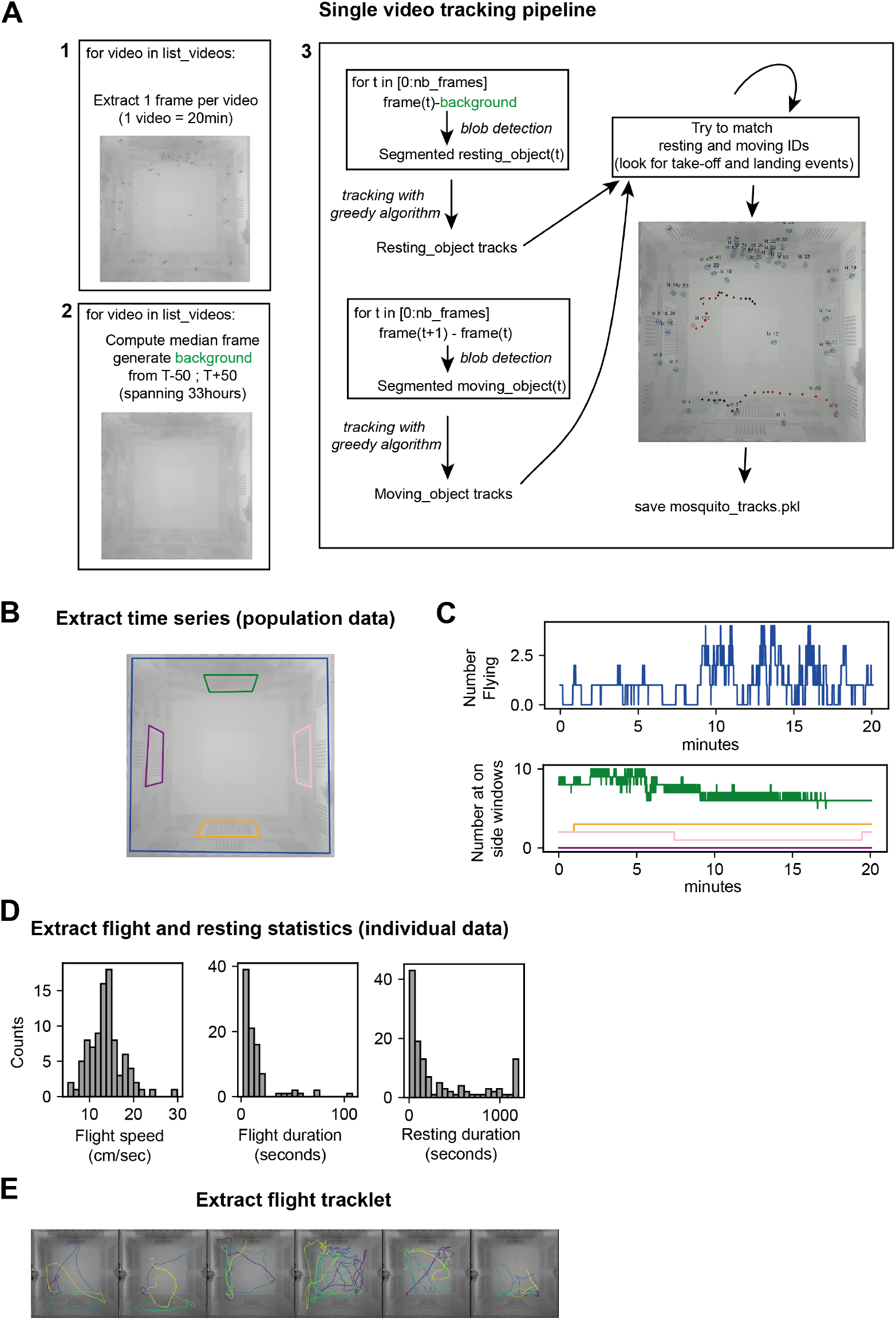
Single video analysis and mosquito tracking. **A**. Our analysis pipeline utilizes standard computer vision methods to segment mosquitoes from the cage background, employing techniques such as background subtraction and thresholding. Accurateness of this approach relies on obtaining a clean background image devoid of mosquitoes but closely resembling the videos with mosquitoes. We achieve this by averaging frames from multiple videos spanning over a day. While mosquitoes can rest in the same spot for several hours, over a 24-hour period, they will move at least once. By averaging 100 frames from 100 consecutive 20-minute videos (one frame per 20 minutes over 30 hours, step 1), we obtain a background image without mosquitoes (step 2). We then perform two distinct operations in parallel to obtain tracks of resting and flying mosquitoes. This separation is based on the observation that, with our camera’s frame rate (25 FPS) and resolution, flying mosquitoes appear relatively blurred and low contrast compared to the sharp, dark appearance of resting mosquitoes. Therefore, tracking them separately improves overall performance. For resting mosquitoes, we subtract the background from all frames and apply a blob detection method to segment and track mosquitoes using their centroids. A greedy algorithm is used on these centroids to obtain tracks of resting mosquitoes, which include those not moving or moving slowly by walking. For flying mosquitoes, instead of using the background image, we subtract the previous frame, as is standard for motion detection in videos. We then apply similar segmentation and centroid tracking with parameters optimized for flying mosquitoes. After obtaining tracks for both resting (immobile or walking) and moving (flying) mosquitoes, we use a custom algorithm to link these tracks if they belong to the same mosquito. This algorithm detects take-off events when a resting track disappears and a moving track appears. By assuming that for a real take-off event, the end of the resting track and the start of the moving track should be close, the algorithm performs a local search to find the best match. It uses a similar strategy for landing events, linking the end of a moving track with the start of a resting track. The algorithm also attempts to connect different moving tracks. This bootstrapping procedure is iterated five times, logging the percentage of matched or unmatched tracks. While imperfect, it does not affect the robustness of population-level variables, such as the fraction of mosquitoes flying or resting on a given surface like the sugar feeder. The final mosquito tracks are stored as a single pickle file for each video analyzed. These files contain the coordinates of 40 mosquitoes at 25 FPS resolution for 20 minutes (including corresponding IDs) and typically range from 20-30 MB in size. **B**. In the GUI app, we can manually draw the different region of interest for analysis: the exact border of the cage (any detection outside of this boundary will not be taken into account) and the 4 windows, behind which there is sugar cotton for instance. In that example, the green frame indicates the boundary of the sugar feeder window, orange the opposite control area, and the purple and pink to other control areas. During mosquito tracking, the number of resting mosquitoes within each window is registered over time. **C**. Example of time series (population level variable) extracted for a single video based on the algorithm in A and boundaries defined in B. **D**. Example of histogram (individual track variable) of flight speed, flight duration and resting time for the entire video. For resting times, the last bin on the right at 1200 seconds indicates mosquitos that did not move at all for the entire video. **E**.Example flight trajectories extracted from the same video, color indicates time, normalized to the duration of each flight trajectory.

**Figure S3.**
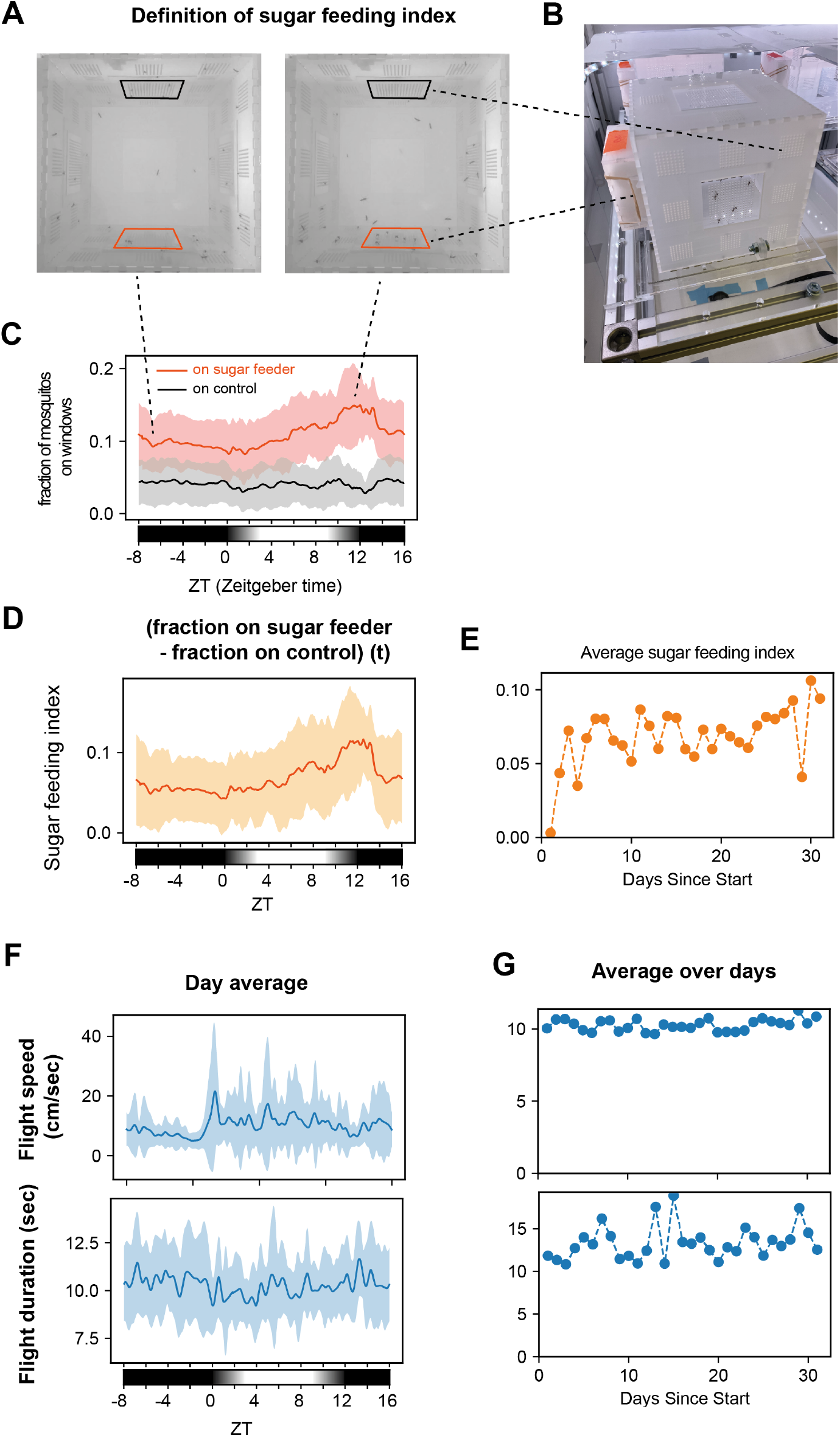
sugar feeding and other flight variables. **A**. Snapshot from movie recorded in Fig.1 experiment during the night (0h30 left) and at the evening peak (19h30 right). Areas manually selected in the app for sugar feeder (orange) and control (black) are overlayed. **B**. Picture of a BuzzWatch cage being monitored with 40 mosquitoes inside, sugar feeder filled with soaked cotton on the left and nothing on the 3 other side windows. **C**. Day average profile (from 32 days) of fraction mosquitoes resting on the sugar feeder window (orange), or the control opposite window (black). Lines indicate the average over 32 days and shaded area are the s.t.d. over 32 days. 20-minutes rolling average was used before averaging different days. **D**. Day average profile of the difference between fraction mosquitoes resting within the sugar feeder window and the fraction resting on the control window opposite. We aimed at quantifying sugar-feeding behavior. Due to the optical setup’s limitations, which prevent direct visualization of proboscis contact with the sugar meal, we implemented an alternative strategy to estimate the number of mosquitoes feeding on sugar at a given time. In the cage, mosquitoes have access to a cotton patch soaked with sugar water through a square window in the sidewall (A-B). We analyzed tracking data to determine the fraction of mosquitoes resting on this sugar-feeder window (orange square in A). and compare it to a control window on the opposite side (black square in A), which has similar geometry but no sugar cotton behind it. By calculating the daily averages, we found that a significantly higher fraction of mosquitoes rested on the sugar-feeder window (orange line C) compared to the control (black line C). This difference was most pronounced around ZT12 (start of dark period), coinciding with the peak of evening flight activity (see Fig.1). These findings suggest that sugar-feeding activity can be indirectly detected by comparing these two curves. For simplicity, we used the difference between these two curves as a “sugar feeding index” (orange line in D). **E**. Sugar feeding activity averaged per day over the course of the 32 day experiment from Fig.1. **F**. Daily average of flight speed (top) and flight duration (bottom) over the 32 day experiment from Fig.1. **G**. Average for each entire day over the course of the 32 day experiment from Fig. 1 for flight speed (top) and flight duration (bottom). Regarding flight duration and speed averaged over days, we did not observe any clear trends over the 32 days of the experiment.

**Figure S4.**
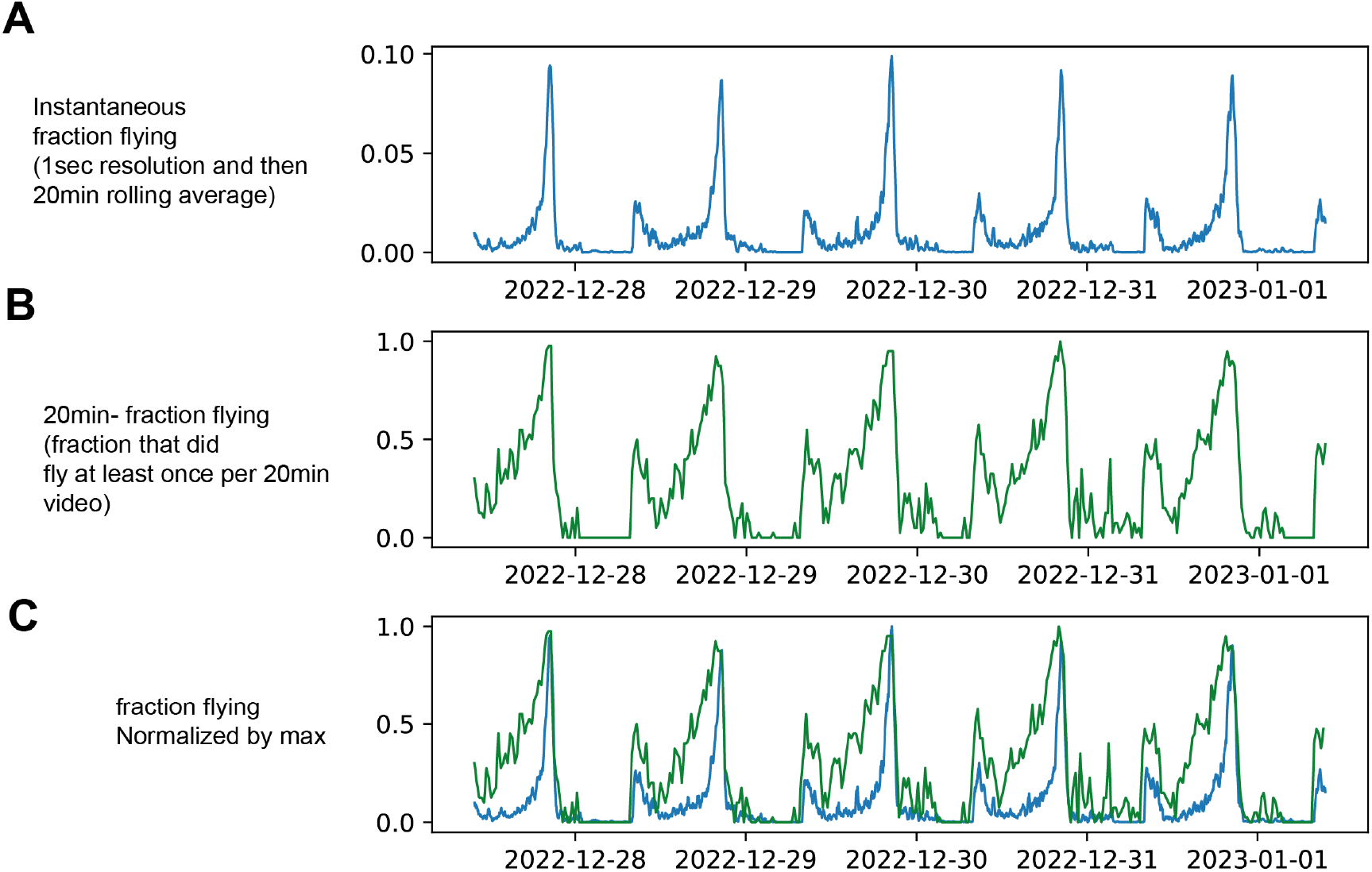
Instantaneous vs global fraction flying. Example data over 5 days from Fig.1. **A**. “Instantaneous” fraction of mosquitoes flying over 5 days. Initially, the time series represents the number of flying mosquitoes at 1-second resolution. The data is resampled to 1-minute intervals (averaged), smoothed with a rolling 20-minute window average, and normalized by the total number of alive mosquitoes (typically 40, decreasing if mosquitoes die during the experiment). **B**. “Global” fraction flying over 5 days. For each 20-minute interval, we extract the distribution of resting times from individual mosquito tracks. To estimate the number of mosquitoes that did not fly during the entire video, we calculate the number of tracks with resting times of 1200 seconds (the 20-minute video duration). We derive the same variable X for each 20-minute video and report (Number_alive_mosquitoes − *X*)*/*Number_alive_mosquitoes (green curve). This provides the fraction of mosquitoes that flew at least once within a 20-minute window. **C**. “Instantaneous fraction flying” and “20-minute global fraction flying” are normalized by their maximum value over the 5-day window considered here. Interestingly, at the peak of mosquito activity, only about 10% of the population is flying at any moment, indicating that most mosquitoes spend their time resting (A). However, when we measured the total number of mosquitoes that flew at least once during a 20-minute period, the number reached nearly 100% during the evening peak (B). This suggests that all mosquitoes take flight eventually in the evening. Due to individual differences in flight timing and their short flight duration (10-20 seconds), the percentage of mosquitoes flying at any given moment is only around 10-20% at most. While these two measurements of flight activity provide slightly different perspectives on flight behavior, they largely overlap (C). For the remainder of the study, we chose to focus on the instantaneous fraction flying, as it is more robust to quantify and more directly linked to the transition rate between resting and flying.

**Figure S5.**
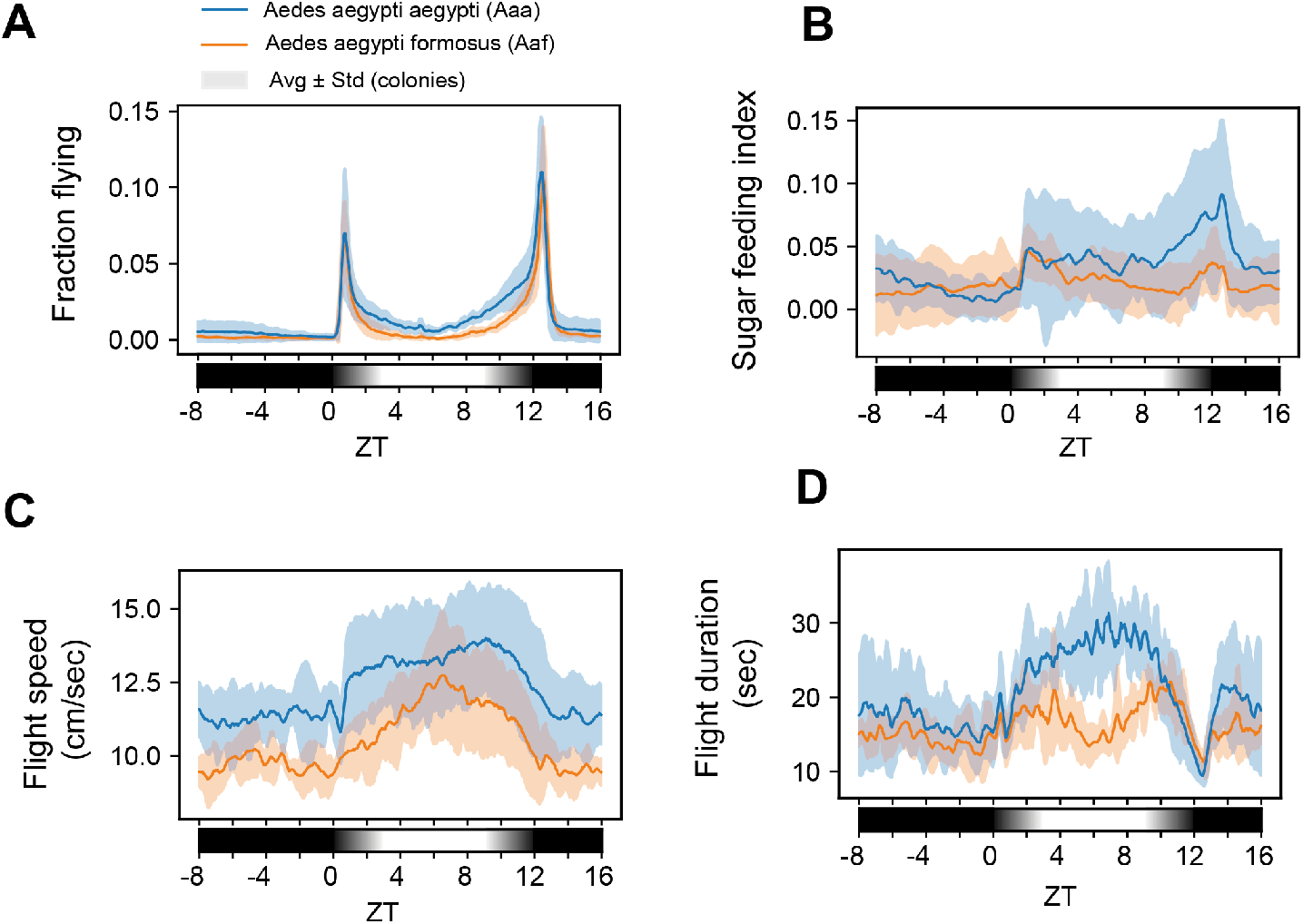
Day-averaged variable comparison between Aaa and Aaf colonies. Average over 5 colonies for Aedes aegypti aegypti (blue) and Aedes aegypti formosus (orange), shaded area is std. over colonies. 20 minutes – moving average; average over 10 days for each colony. **A**. fraction flying **B**. Sugar feeding index **C**. Flight speed **D**. Flight duration

**Figure S6.**
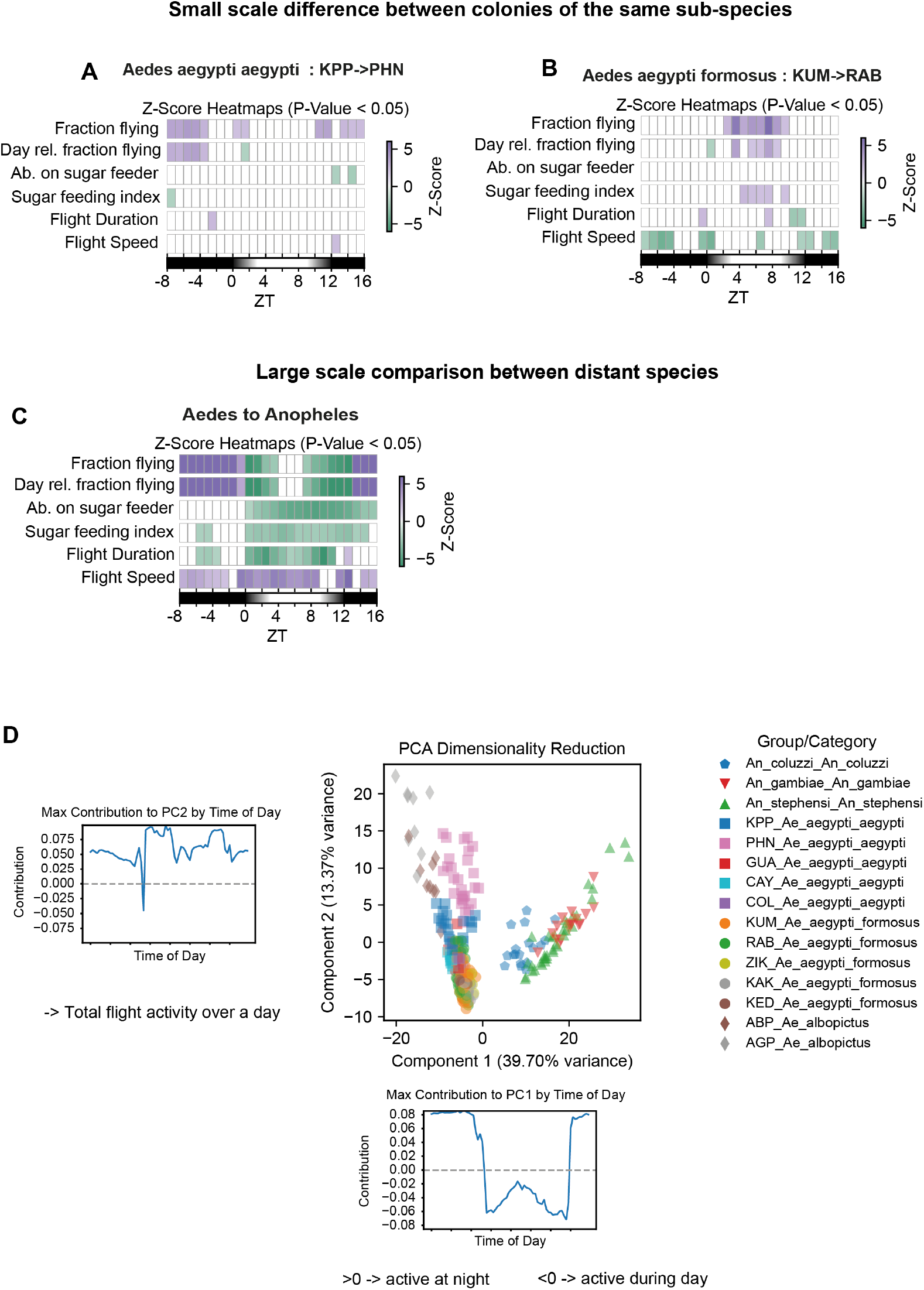
Species and colony level analysis. **A**. Barcode GLMM plot between two Aedes aegypti aegypti colonies : KPP (Thailand) colony and PHN (Cambodia) colony. Each colony was measured with 3 independent replicate cages over 10 days (40 females). KPP colony is used as baseline. **B**. Barcode GLMM plot between two Aedes aegypti formosus colonies : KUM (Ghana) colony and RAB (Kenya) colony. Each colony was measured with 3 independent replicate cages over 10 days (40 females). KUM colony is used as baseline. **C**. Barcode GLMM plot between Aedes (Aedes aegypti and Aedes albopictus) and Anopheles species (gambiae, stephensi, coluzzii). Aedes is taken as baseline. **D**. Pattern based analysis for all Aedes and Anopheles fraction flying data. Each dot is a single day of measurements. Every cage was monitored for at least 10 days, only Ae. aegypti colonies KPP,PHN,RAB and KUM were monitored in three independent replicate cages. On the left, hour based contribution to PC2 and on the bottom, PC1. After focusing on the difference between Ae. aegypti sub-species, we wanted to enlarge the scope and see how our methods would behave when e.g. investigating differences between species and colonies. Using our local GLMM approach, we compared two different colonies for each sub-species of Ae. aegypti (KUM and RAB for Aaf and KPP PHN for Aaa) and found statistical differences for the fraction flying over time, but less clearly for other behavioral variables. For instance the Aaa colony from Phnom Penh (PHN) happened to be more active at night compared to the Thailand strain (KPP) (A); while the Aaf colony from Kenya (RAB) was more active between 11h-17h than the colony from Ghana (KUM) (B). These specific differences between colonies were of similar magnitude within the two sub-species, and could reflect specific adaptation to an ecological niche or latitude. At a much larger scale, we compare different between Anopheles, known to be active at night and Aedes, known to be active during the day. The local ‘barcode” analysis revealed strong differences for all variables at almost all times of day, as we could expect from such different mosquito species. We then also used our global dimensionality reduction approach to compare the pattern of 3 different Anopheles species, the two Aedes sub-species and Ae. albopictus (D). This approach revealed that on the scale of difference between distant species, the observed distinction between Aaa and Aaf is still visible, but relatively minor compared to difference with Ae. albopictus.

**Figure S7.**
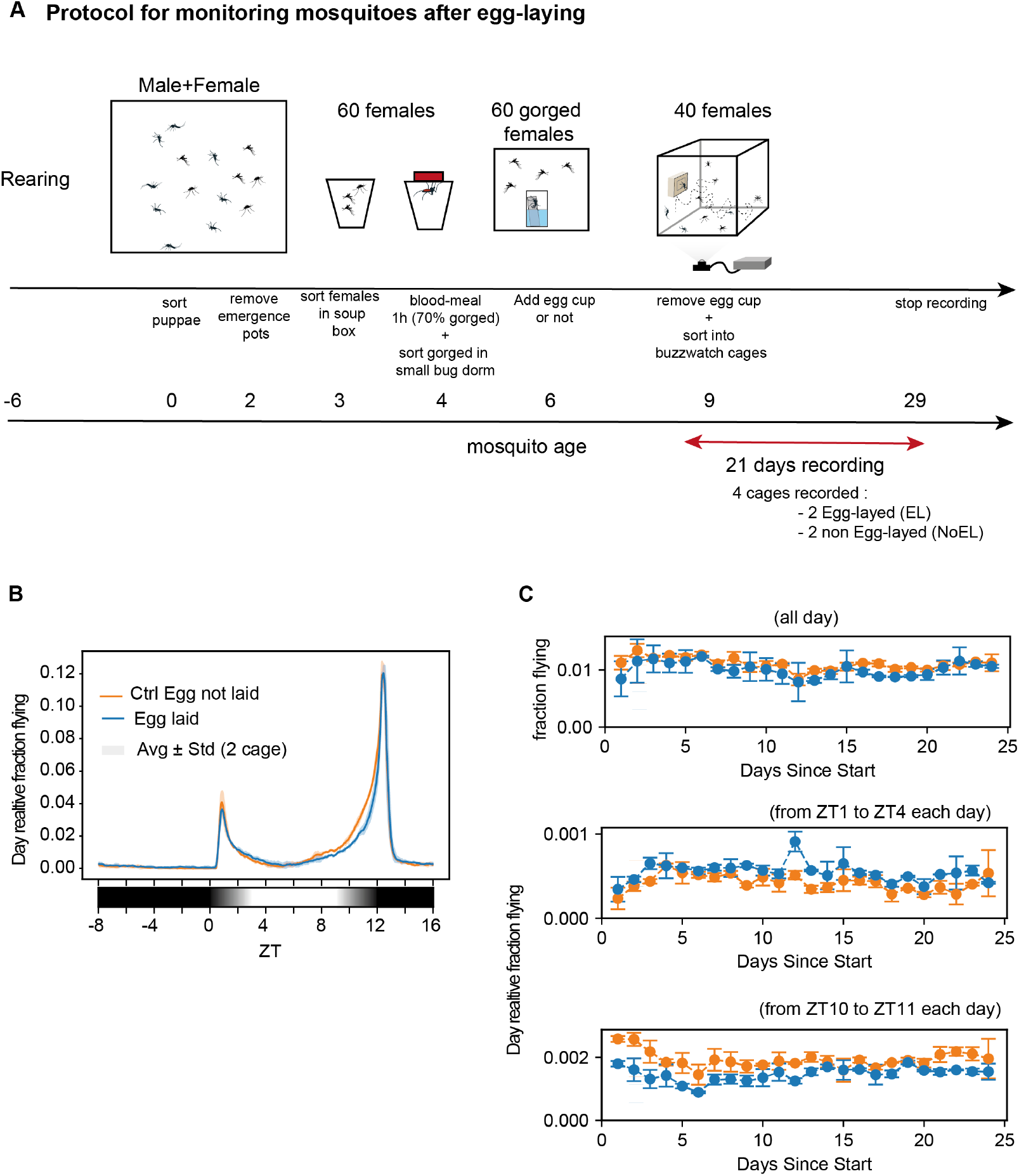
Egg-laying experiment. **A**. Protocol to estimate the effect of egg-laying on mosquito flight behavior. After rearing and mating, females were sorted on ice and allowed to have a blood meal for 1 hour. Fully engorged females were sorted on ice and placed in a box (60 females per box) for 6 days with either a beaker filled with water and Whatman paper to lay eggs, or nothing. Most of the egg laying happens 3-4 days after the blood meal. Females were then sorted on ice and transferred to BuzzWatch cages, 40 females per cage and 2 cages per condition (egg laid versus not laid). Mosquito behavior was monitored over 21 days, sugar was changed weekly. **B**. Averaged day-relative fraction flying (fraction flying normalized by cumulative sum of fraction flying over each day) for mosquito that laid eggs (blue line) or did not lay eggs (orange line). Day relative time series was averaged from 24 days, then average between 2 replicate cage. Shaded area corresponds to standard variation over 2 replicate cages. **C**. Average fraction flying per day for mosquito that laid eggs (blue) or did not lay eggs (orange). (Top) “absolute” fraction flying averaged over entire days for each day. (Middle) Day-relative fraction flying averaged between ZT1 and ZT5 or ZT10 and ZT11 (Bottom). Error bars are standard deviation between 2 replicate cages.

## Supplementary text - Flight Speed from 2D vs 3D Trajectories

In our study, we analyze 2D flight trajectories of mosquitoes from BuzzWatch video recordings, which are projections of their actual 3D trajectories. This leads to an underestimation of the average flight speed as the displacement in the *z*-direction (perpendicular to the camera’s focal plane) is unaccounted for.

Assuming isotropic flight, where flight speed is uniformly distributed across the *x, y*, and *z* axes, we estimate that the 2D measured speed *v*_2*D*_ is related to the actual 3D speed *v*_3*D*_ by:

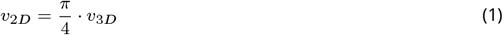

This conversion factor of 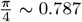 derives from the geometric projection of a 3D vector onto a plane. To see that, let us consider a unit vector **v** in spherical coordinates with *θ* the polar angle and *ϕ* the azimuthal angle (https://en.wikipedia.org/wiki/Spherical_coordinate_system) :

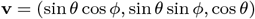

When projected onto the *xy*-plane, the *z*-component is ignored, yielding:

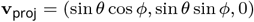

The magnitude of this projection is:

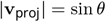

To find the average projection length over all orientations, we integrate over the unit sphere. The elemental surface area in spherical coordinates is sin *θ dθ dϕ*. Thus, the average projection *L*_avg_ is:

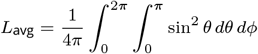

Utilizing [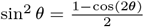, we find:

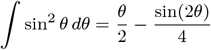

Evaluating from 0 to *π*:

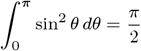

The integration over *ϕ* results in 2*π*, thus:

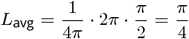

Therefore, the average projection length of a unit vector onto a 2D plane is 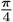, supporting the conversion factor used.

